# The Internal Structure of Metacommunities

**DOI:** 10.1101/2020.07.04.187955

**Authors:** Mathew A. Leibold, F. Javiera Rudolph, F. Guillaume Blanchet, Luc De Meester, Dominique Gravel, Florian Hartig, Pedro Peres-Neto, Lauren Shoemaker, Jonathan M. Chase

**Author notes:** Data accessibility statement:* Should this manuscript be accepted, the data and code supporting the results will be archived in an appropriate public repository such as Github and will be mirrored at Zenodo or similar locations.

## Abstract

Current analyses of metacommunity data largely focus on global attributes across the entire metacommunity, such as mean alpha, beta, and gamma diversity, as well as the partitioning of compositional variation into single estimates of contributions of space and environmental effects and, more recently, possible contributions of species interactions. However, this view neglects the fact that different species and sites in the landscape can vary widely in how they contribute to these metacommunity-wide attributes. We argue for a new conceptual framework with matched analytics with the goals of studying the complex and interactive relations between process and pattern in metacommunities that is focused on the variation among species and among sites which we call the ‘internal structure’ of the metacommunity. To demonstrate how the internal structure could be studied, we create synthetic data using a process-based colonization-extinction metacommunity model. We then use Joint Species Distribution Models to estimate how the contributions of space, environment and biotic interactions driving metacommunity assembly differ among species and sites. We find that this approach to the internal structure of metacommunities provides useful information about the distinct ways that different species and different sites contribute to metacommunity structure. Although it has limitations, our work points at a more general approach to understand how other possible complexities might affect internal structure and might thus be incorporated into a more cohesive metacommunity theory.

## Introduction

Community ecology is currently undergoing an important renaissance in both its concepts and tools. One of the more exciting and important elements of this renaissance is in the use of the metacommunity concept, which recognizes the feedback between local communities and the broader-scale regional biota or species pools (Hanski and Gilpin 1991, Leibold et al. 2004 and reviewed in Leibold and Chase 2017). Early metacommunity studies tended to focus on specific scenarios that involve such feedbacks (e.g., Levins and Culver 1971, Horn and MacArthur 1974, Levin 1974, Sloan-Wilson 1992, Leibold 1998, Hubble 2001, Amarasekare and Nisbet 2001). These were later synthesized into several (discrete) categories of metacommunities dynamics (Leibold et al. 2004). While these categories proved useful, it is now apparent that there is a much more complex and nuanced spectrum of possibilities regarding the mechanisms and processes underlying the structure of metacommunities (see Leibold and Chase 2017). Ongoing developments, including both more sophisticated theoretical (e.g., Shoemaker and Melbourne 2016, Fournier et al. 2017, Ovaskainen et al. 2019, Thompson et al. 2020) and analytical (e.g., Legendre and De Cáceres 2013, Hui et al. 2013, Ovaskainen et al. 2017, Ohlman et al. 2018, Jabot et al. 2020) approaches, aim for a deeper understanding of the regional-local community-level feedbacks. Understanding these more subtle feedbacks between local communities and the regional biota also has important implications that extend to applied ecology, as well as environmental and health concerns (e.g., Bengtsson 2009, Schiesari et al. 2019, Miller et al. 2019, Brown and Barney 2020).

Despite the progress we observed in the study of metacommunities, two issues remain central in contemporary metacommunity analyses. The first is that most theoretical frameworks operate under the assumption that processes act similarly on all species and sites so that it makes sense to infer, for example, that an entire metacommunity being dominated by neutral processes or species sorting. A second limitation is that the most widely used analytical frameworks in metacommunity ecology assume that community assembly is dominated by spatial and environmental factors, without considering the influence of biotic interactions. In reality, however, variation among species and among sites, as well as biotic interactions, can interact in complex ways to produce metacommunity patterns. For instance, the spatial structure of environmental features can vary within landscapes (i.e., among sites) which, in turn, can affect the ways species interact and are sorted into local communities (Peres-Neto et al. 2012). Thus, in contrast with many previous studies (e.g., Blanchard et al. 2020, Jabot et al. 2020), we emphasize that species can be heterogeneous in how they contribute to metacommunity level properties and that different sites can also vary in how they contribute to these patterns (as suggested by earlier work by e.g. Pandit et al. 2009, Legendre and De Cáceres 2013).

If we acknowledge that community assembly within a metacommunity is a complex process that involves heterogeneous contributions of species sorting, interactions, dispersal, and stochasticity acting on a regionally-defined species pool within a given landscape (e.g. Vellend 2010, 2016, Weiher et al. 2011), the question becomes - how can we sensibly document these processes from observational data in a way that is useful for ecological understanding? Current analytical frameworks for analyzing metacommunity assembly, such as diversity metrics, coexistence patterns and variation partitioning analysis (e.g. Borcard et al. 1992, Gotelli and McCabe 2002, Leibold and Mikkelson 2002), describe global (i.e. mean) metacommunity properties. Recent efforts have used several of these global metrics to dissect the relative importance of different major classes of metacommunity processes (Ovaskainen et al. 2019, Guzman et al. 2021). While these approaches provide insights into the processes that drive species distributions and determine their levels of interaction within metacommunities, they only characterize global (general) attributes across the entire metacommunity, which we consider to be the *external* structure of metacommunities. Here, we focus on the *internal* structure of metacommunities that focuses on the contributions of individual species and individual sites (or patches) to the global (i.e., mean) metacommunity structure (see also Fournier et al. 2017, Suzuki and Economo 2021).

To illustrate the advantages of studying the internal structure of metacommunities, we created synthetic data from a process-based metacommunity model structured by competition-extinction dynamics. Using simulations has the advantage that we know the true underlying processes and therefore we have clear expectations about what we should infer from the generated data.

We then use joint species distribution models to analyze the resulting distribution data (JSDMs; see review in Warton et al. 2015). JSDMs are multivariate regression models that simultaneously describe metacommunity structure as a function of species-specific environmental preferences, spatial autocorrelation, and covariances among species. We have adapted JSDMs to estimate both the contributions per site and per species for these three components of variation. With due caution, one can relate environmental predictors to the fundamental environmental niche of the species, spatial effects to dispersal, and covariation to species interactions. Among the currently available statistical methods, JSDMs arguably extract the greatest amount of variation from spatial community data (Warton et al. 2015, Ovaskainen et al. 2017), even though they have been facing increased scrutiny about the interpretation of estimates and potential bias (e.g. Poggiato et al. 2020, Miele et al. 2021, Blanchet et al. 2020, but see Pichler & Hartig, in press).

By simulating data from a process-based model, we can address the extent to which JSDMs are able to correctly separate environmental, spatial, and biotic effects. It also allows us to explore how we can use their outputs to generate a more detailed and accurate picture of the internal metacommunity structure. The latter is possible because JSDM results can be decomposed (as we show later) into species-specific and site-specific contributions of environment, space and biotic interactions. Our results show that heterogeneity among species attributes can cause substantial variation in their metacommunity patterns, as identified using JSDMs. Some of this heterogeneity can be identified by JSDMs and be related to variation among species in their attributes such as dispersal and similarities (or differences) on their spatial associations with other species; or uniqueness among sites regarding environmental, spatial attributes (e.g., connectivity) and species associations. We show how JSDMs can be used to estimate the piece-wise contribution of species (what we call the ‘internal structure’ of the metacommunity) to evaluate the metacommunity-wide overall effects of environment, space, co-distributions and stochasticity (which we think of as the ‘external’ structure), we can thus evaluate how variation among species affect overall metacommunity structure. We also show that a similar dissection of variation can be made among sites and argue that this can also be thought of as a distinct aspect of the ‘internal structure’ that describe how sites differ in their contributions to overall patterns of metacommunity structure to reveal at least some components of the species-level and site-level contributions to overall metacommunity variation.

## Methods to quantify the link between process and pattern using a simple metacommunity simulation and refined statistical approach

To test and exemplify our ability to infer individual species and site contributions, we simulated data from a process-based metacommunity model, which allowed us to create observations with full knowledge about the underlying mechanisms. Our process-based analytical model is based on a spatial implementation of spatially implicit site occupancy models (e.g., Levins and Culver 1971, Horn and MacArthur 1972, Levin 1974, Hastings 1980, Hanski 1991) to describe dynamics in heterogeneous metacommunities. We focused here on a model for predicting presence-absence (and not abundance; but see Supporting Information) because it is the most widely available type of empirical data for metacommunity analyses. For each species in each patch, we model occupancy using two key equations (details in the Supporting Information). The first of these describes the colonization of patch *z* by species *i* during a discrete time interval, Δ*t*:

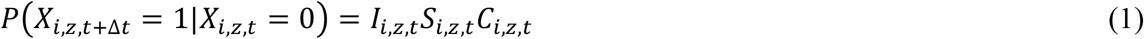

where *X*_*i,z,t*_ is a stochastic variable representing the occurrence of species *i* at location *z* at time *t,I*_*i,z,t*_ is the number of immigrants of species, *S*_*i,z,t*_ is the effect of environmental filtering on the probability of establishing a viable local population, and *C*_*i,z,t*_ is the effect of ecological interactions on the establishment probability. Second, we consider the alternative possibility - the extinction of species *i* in patch *z* during the time interval Δ*t*:

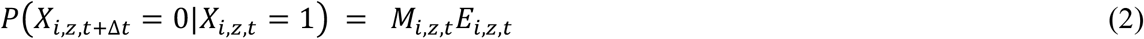

where *M*_*i,z,t*_ and *E*_*i,z,t*_ are the responses of the extinction probability to the local environment and to ecological interactions, respectively.

At steady state the solution to this model is:

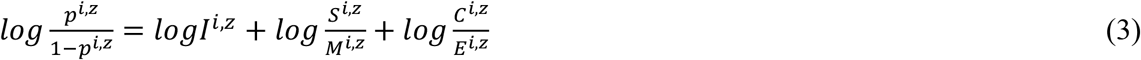

where *p*^*i,z*^ is the expected probability that site *z* is occupied by species *i*. This formulation assumes that immigration(*I*^*i,z*^), ‘environmental selection’ (*S*^*i,z*^ ∧ *M*^*i,z*^) and interactions (*C*^*i,z*^ ∧ *E*^*i,z*^) can be separated into distinct effects.

Equation 3 suggests that distributions of species in a metacommunity can be studied by correlation-based methods such as JSDMs to separate the contributions of these effects into spatial effects (driven by immigration), environmental filtering (driven by abiotic selection) and species co-distribution unrelated to either space or environment, with an additional fraction quantifying residuals resulting from stochasticity in the case of a finite number of patches (see also Shoemaker et al. 2020). Furthermore, the likelihood of every observation can be marginalized over each species (by summing the likelihoods for a given species across all patches) to describe the variation among species. Alternatively, the likelihood can be marginalized by sites (by summing the likelihoods for a patch across all species) to describe the variation across the metacommunity landscape. In doing so, we can quantify the importance of environment, species co-distribution, and space for predicting metacommunity structure as a whole, as well as quantify their importance for predicting the presence-absence (or, in principle, the abundance) of individual species or community composition at individual patches. As JSDM are fundamentally correlational, they can only evaluate the degree to which observations are consistent with our model but (as in most correlative models) cannot be used as definitive tests of causation.

To illustrate the utility of our analytical model in a more realistic framework that also includes stochasticity and spatially explicit landscapes, we implemented the key processes of drift, environmental filtering, dispersal, and species interactions (Vellend 2010, 2016) in a flexible simulation version of the model above (described in more details in the Supporting Information). The simulation model allows us to vary each process separately for each species in a heterogeneous spatially explicit landscape. It simulates the dynamics of a metacommunity across a set of patches and generates a spatial network that specifies the connectivity among patches. The state variables of the simulation are the occupancy of each species in every patch (i.e. presence/absence, though future implementations could also address abundance data, e.g. Rybicki et al. 2018, Ovaskainen et al. 2019, Thompson et al. 2020). Each patch can be colonized from nearby patches depending on their location in the landscape, dispersal rate of the species and proximity of extant populations in neighboring patches. Each species in each patch is subject to extinctions that reflect demographic and/or environmental stochasticity. Patches can differ in local environmental conditions that differentially influence baseline colonization and extinction probabilities. Species interactions are modeled in two ways. First, the presence of other species in a patch can modify baseline colonization probability (a reduction in the case of competition). Second, co-occurring species can modify baseline extinction probability (an increase in the case of competition).

We next apply a JSDM to the resulting distribution of species among patches. Specifically, we use the HMSC R package, a modified version of the HMSC package described by Ovaskainen et al. (2017). The new version of HMSC has been modified to incorporate Type III sum of squares errors and site-by-site variation decomposition into variation partitioning analysis (Blanchet, 2019). With this implementation, we model species distributions as a function of the environment, spatial autocorrelation and species co-distributions (see Supporting Information for technical details). After fitting the model using HMSC, we partition variation in the distribution of species in the metacommunity into four statistical components (or fractions) using an approach akin to classic variation partitioning (Borcard et al. 1992; Peres-Neto et al. 2006). Details about how this variation partitioning is computed through HMSC is given in the Supporting Information. In particular, we simplified the 8-way resulting variation components into 4 more easily visualized components to quantify the effects of environment (labeled [E]), spatial patterning (labeled [S]), co-distribution among species (labeled [C]) and residual (unexplained) variation that cannot be attributed to any of the three previously mentioned fractions (i.e. sets of predictors). The latter is expressed as 1-R^2^, where R^2^ is the proportion of variation explained by the model and includes fractions [E], [S], and [C]. Aggregating independent and non-independent fractions as explained in the Supplementary Information results in a loss of more detailed information but allows us to visualize and simplify our interpretation of the results in ways that are useful for the present study.

Although the analytical model described in equations 1-3 suggests that making links between processes and patterns using JSDMs are possible, we wished to evaluate if this was also likely in less idealized situations such as those used in our metacommunity model. We thus simulated a number of scenarios (i.e., ‘thought experiments’) that vary the strength of environmental selection, dispersal and competition. Comparing scenarios with varying niche breadth and competition (scenarios A-D, Fig. 2 and Fig 3), and a more complex case where species compete and vary in both dispersal and in their responses to the environment (scenario G, Figure 4, scenarios E and F in the SI) highlight how HMSC provides an avenue for distinguishing between underlying processes based on abundances of species across metacommunity patches. Using our framework, our goal here was to illustrate how links between pattern and process might be inferred in metacommunities under our “internal structure” framework. In doing so, we leave a more extensive and systematic evaluation of the model’s components (e.g., performance of JSDMs under multiple complex scenarios) for future work (but see Ovaskainen and Nerea 2020).

**Figure 1:**
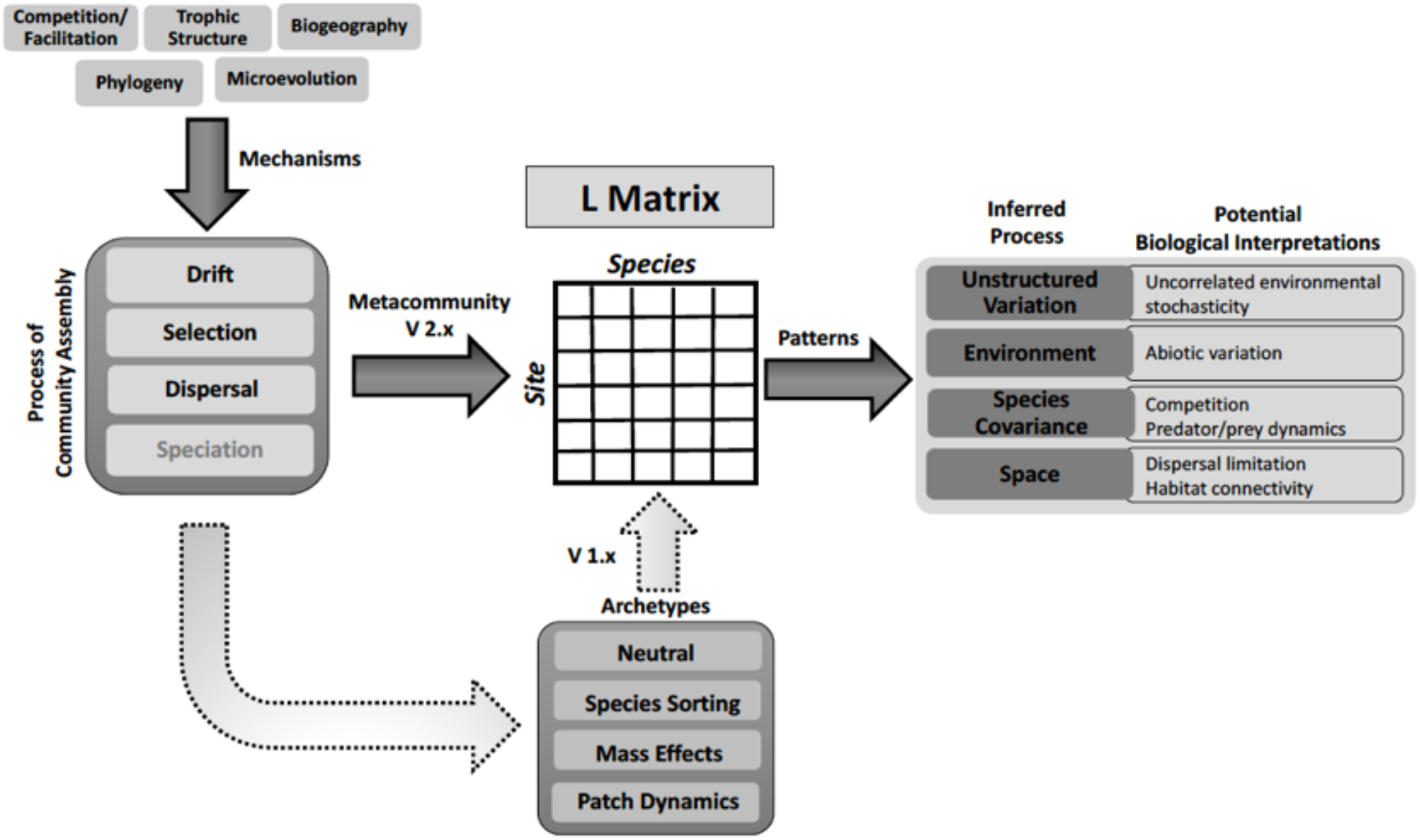
A summary of the metacommunity problem. Species distributions, denoted by the species-by-sites **L** matrix, are the outcome of drift, selection, dispersal, and speciation. These basic processes can be influenced by species interactions, food web structure, biogeography, phylogeny and micro-evolution. Metacommunity theory mainly focuses on drift, selection and dispersal. We view previous approaches based on the four archetypes of Leibold et al. (2004) as being much more indirect and idealized. Instead, we call for a more direct evaluation of how the basic processes affect the **L** matrix, and how to dissect the consequences to the distributions of different species and the occupancy of different sites, for example by using a JSDM to identify main effects and interspecific variability in the importance of unstructured, biotic, environmental, and spatial effects on **L**. This approach allows us to recognize and address the effects of heterogeneities among species and among patches on the overall structure of the metacommunity.

**Figure 2:**
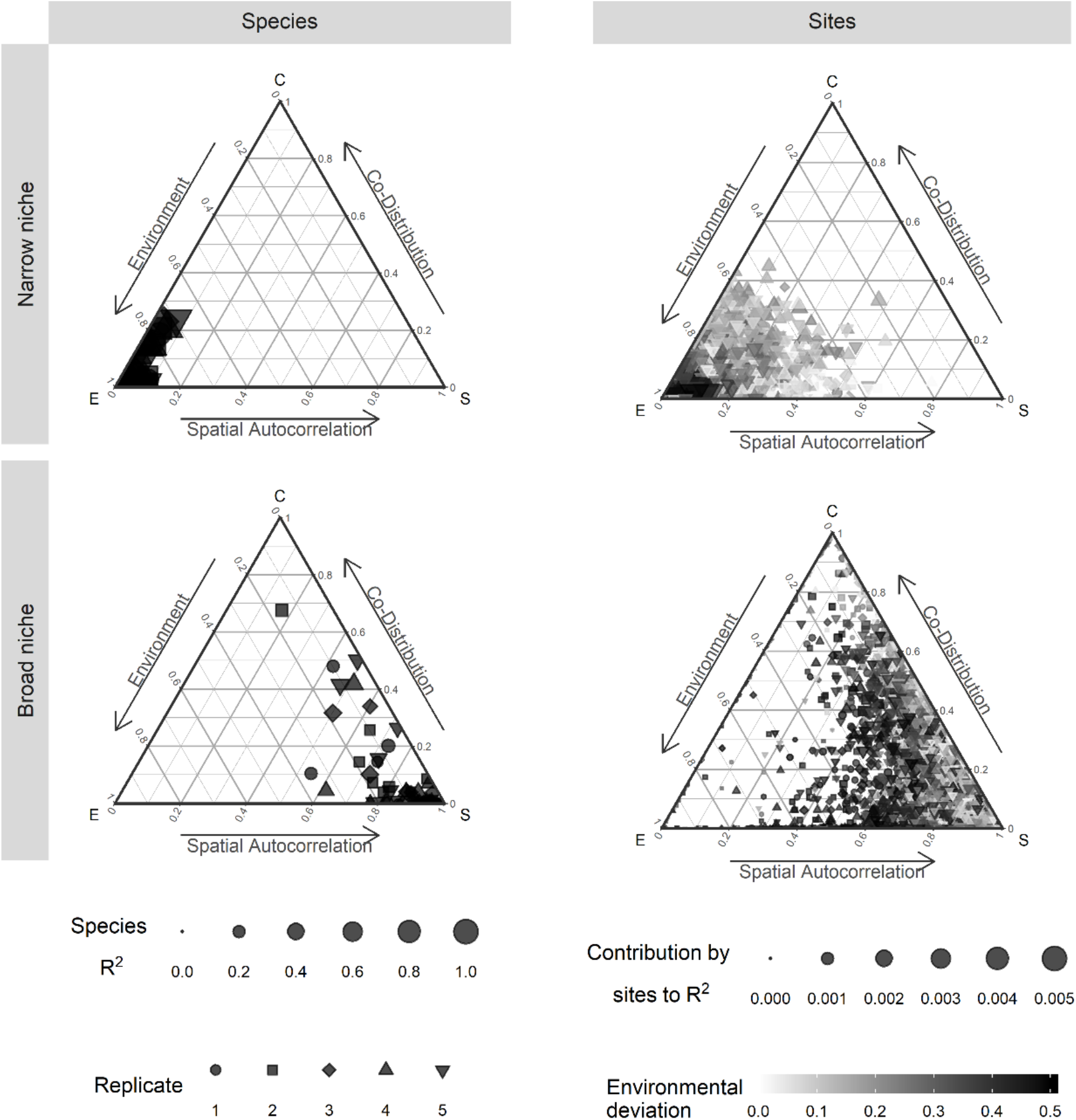
Ternary plots describing the three components of metacommunity internal structure for two different simulation scenarios with no species interactions (independent metapopulations): The upper panels correspond to narrow environmental niches whereas lower panels correspond to wide environmental niches. Each dot represents a species (left panels) or a site (right panels). The size of the symbol is proportional to the R^2^ of the model (note the different scales used for species and sites) and the location indicates the proportion of explained variation attributed to environmental factors (*E -* lower left), spatial effects (*S* - lower right) and remaining co-distributions (*C* - upper apex) (see SI for details). In the species panels (left side) different symbols indicate different replicate simulations; generally, these indicate that the distribution of species responses are variable within replicates but that the overall variation among replicates are repeatable. In the site panels (right side), the shading indicates how central (lighter) or extreme (darker) the local environmental conditions are on the gradient; these also show substantial variation but indicate that more extreme environmental conditions increase the effects of local environment on occupancy patterns than more central conditions.

**Figure 3:**
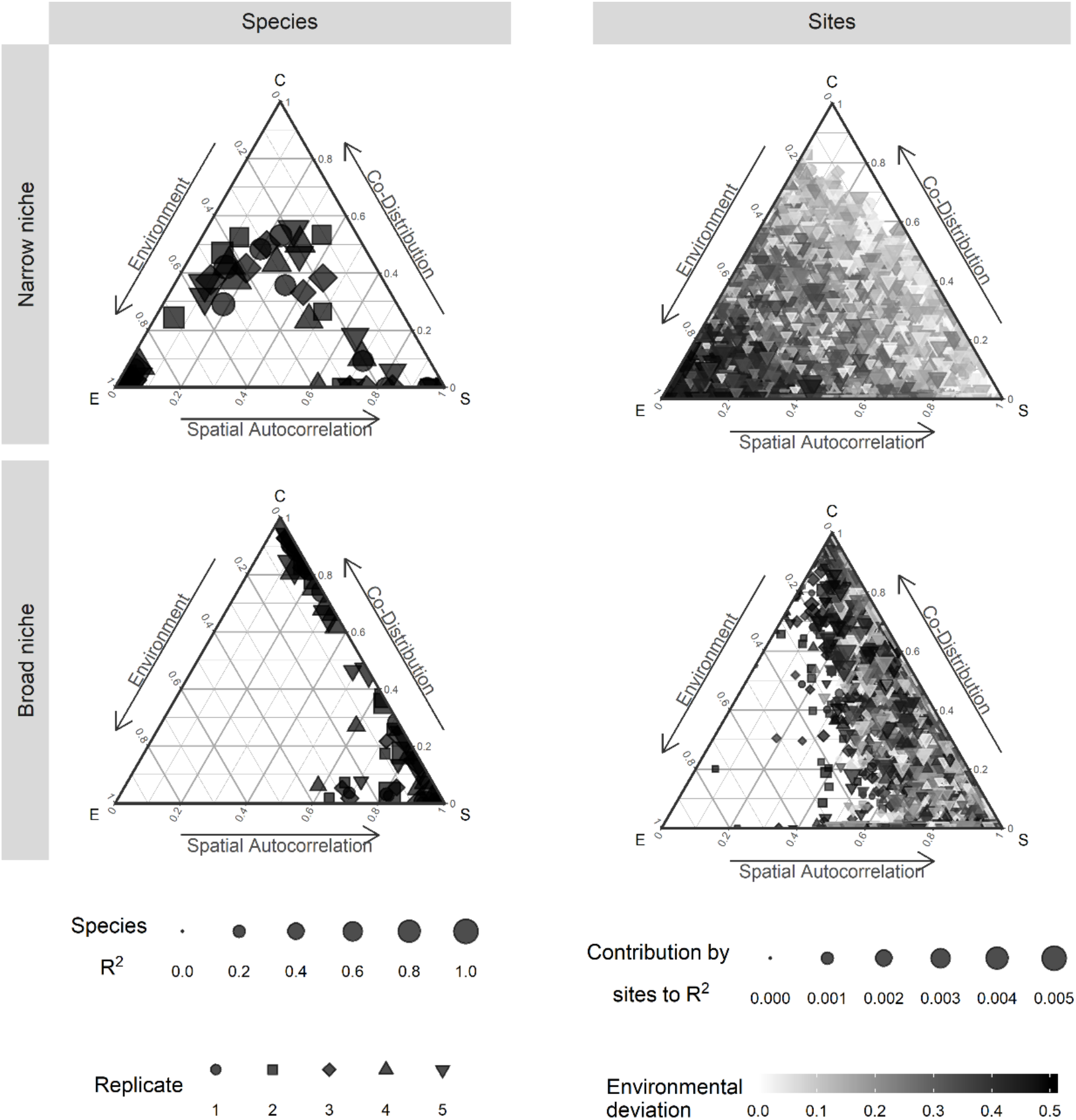
Ternary plots describing the three components of metacommunity internal structure for different simulation scenarios with competition among the species. Notation is the same as in Figure 2. The upper panels correspond to narrow environmental niches whereas the lower panels correspond to wide environmental niches. Left-hand panels show variation components for different species whereas panels on the right-hand side of the figure correspond to variation components for different sites.

**Figure 4:**
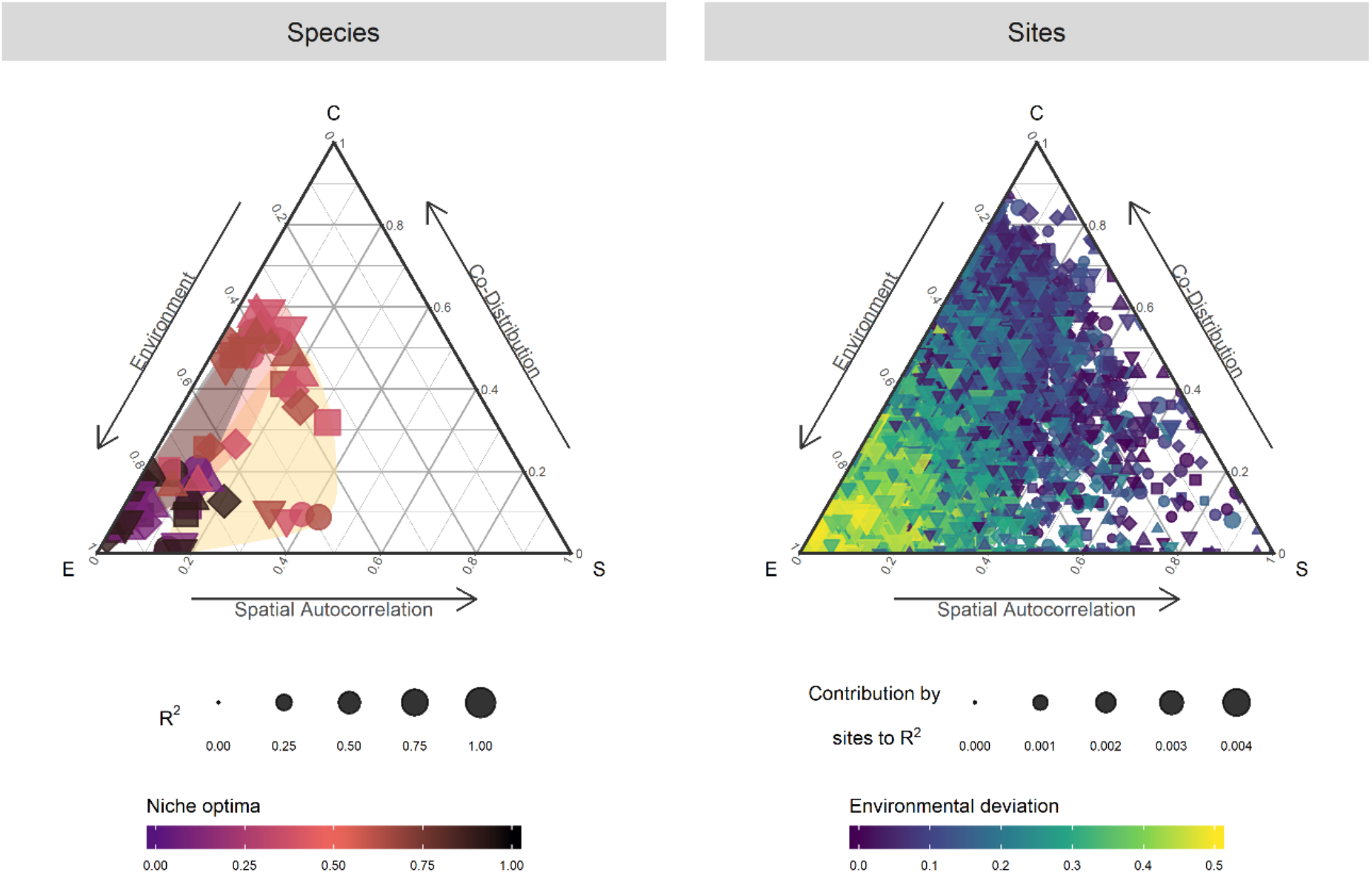
Ternary plots for species (left panel) and sites (right panel) for simulations with species that differ in environmental position along the gradient and dispersal ability. The size of the symbol indicates the R^2^ of the model for each species or site). In the left panel (species) the color indicates the preferred local environmental conditions for species (yellow for species that prefer centrally located environmental conditions, purple or magenta for species with more extreme environmental optima). The symbol indicates the dispersal rate of the species (circles are more dispersal limited, squares are least dispersal limited and triangles are intermediate). In the right-hand panel the color indicates the degree of deviation from centrality along the environmental gradient (as in Figure 2).

## Results of simulation experiments

In a first set of simulations, we considered a situation where species had distinct environmental optima along an environmental gradient and had limited dispersal (Figure 2, Table 1). We contrasted the case where the environmental niches were narrow (steep changes in baseline colonization success and extinction rates with small deviations in environment) with the case with identical optima, but with wide environmental niches (much weaker changes in colonization and extinction with environmental value). As expected, we found that these differences in environmental niche breadth had strong effects on the relative importance of environmental filtering (fraction [E]) versus spatial patterning (fraction [S]). Specifically, we found stronger spatial effects when niche breadths were broad and stronger environmental filtering when niches were narrower (Figure 2, Table 1). We also found that the R^2^ values were higher for the case with narrow niches than with wide niches. Finally, we found non-zero (though relatively weak) variation components for co-distributions (fraction [C]) in both cases, especially when niches were broad even though our analytical model would predict the absence of such variation components since colonization and extinctions were not affected by species interactions in these simulations.

**Table 1:** Summary of metacommunity level variation components for the seven different scenarios modeled in this study.

| Scenario | Corresponding Figure | Fractions |  |  |  |
| --- | --- | --- | --- | --- | --- |
|  |  | E (SD) | S (SD) | C (SD) | Residuals 1-R<sup>2</sup> (SD) |
| A | 2, upper panels | 0.75 (0.11) | 0.026 (0.019) | 0.044 (0.062) | 0.18 (0.11) |
| B | 2, lower panels | 0.019 (0.015) | 0.15 (0.037) | 0.019 (0.036) | 0.81 (0.04) |
| C | 3, upper panels | 0.42 (0.31) | 0.21 (0.2) | 0.14 (0.16) | 0.23 (0.12) |
| D | 3, lower panels | 0.018 (0.026) | 0.35 (0.3) | 0.28 (0.33) | 0.35 (0.33) |
| E | Supplement | 0.63 (0.21) | 0.067 (0.069) | 0.13 (0.15) | 0.18 (0.089) |
| F | Supplement | 0.74 (0.097) | 0.035 (0.025) | 0.047 (0.057) | 0.18 (0.097) |
| G | 4 | 0.5 (0.17) | 0.08 (0.072) | 0.2 (0.16) | 0.22 (0.12) |

We next simulated metacommunities with identical parameters as above, except with added interspecific competition effects (Figure 3, Table 1). As in the case without species interactions (compare with Figure 2), narrow niches enhanced the relative strength of environmental filtering (fraction [E]) and reduced spatial patterning (fraction [S]) when compared to wide niches. In these simulations, however, the co-distribution components (fraction [C]) were much more substantial than without species interactions. We also found that adding interspecific competition substantially increased the total amount of variation explained (R^2^) by the model (i.e. due to the joint component of co-distribution).

We conducted a number of other simulations to explore if interspecific variation on environment (fraction [E]), space (fraction [S]), and co-distributions (fraction [C]) depend on dispersal, niche breadth, and interactions. Illustrative examples are shown in the supplemental information and summary statistics are shown in Table 1. In Figure 4, we present the results from one of these examples that includes heterogenous dispersal to show how the internal structure can reveal how dispersal variation affects species distributions. We found that one could distinguish species by the degree to which their distributions are related to environment (fraction [E]), space (fraction [S]) and co-distributions (fraction [C]) (Figure 5a) and we found that this could be related to their traits (i.e., species optima in our simulation framework). Species with higher dispersal ability and more specialized environmental niche positions had distributions better predicted by the environment than those that were dispersal limited and had distributions that presented a higher level of spatial autocorrelation (fraction [S]). Species with optima closer to the middle of the environmental gradient also had a larger fraction [C] than those with more extreme optima.

**Figure 5:**
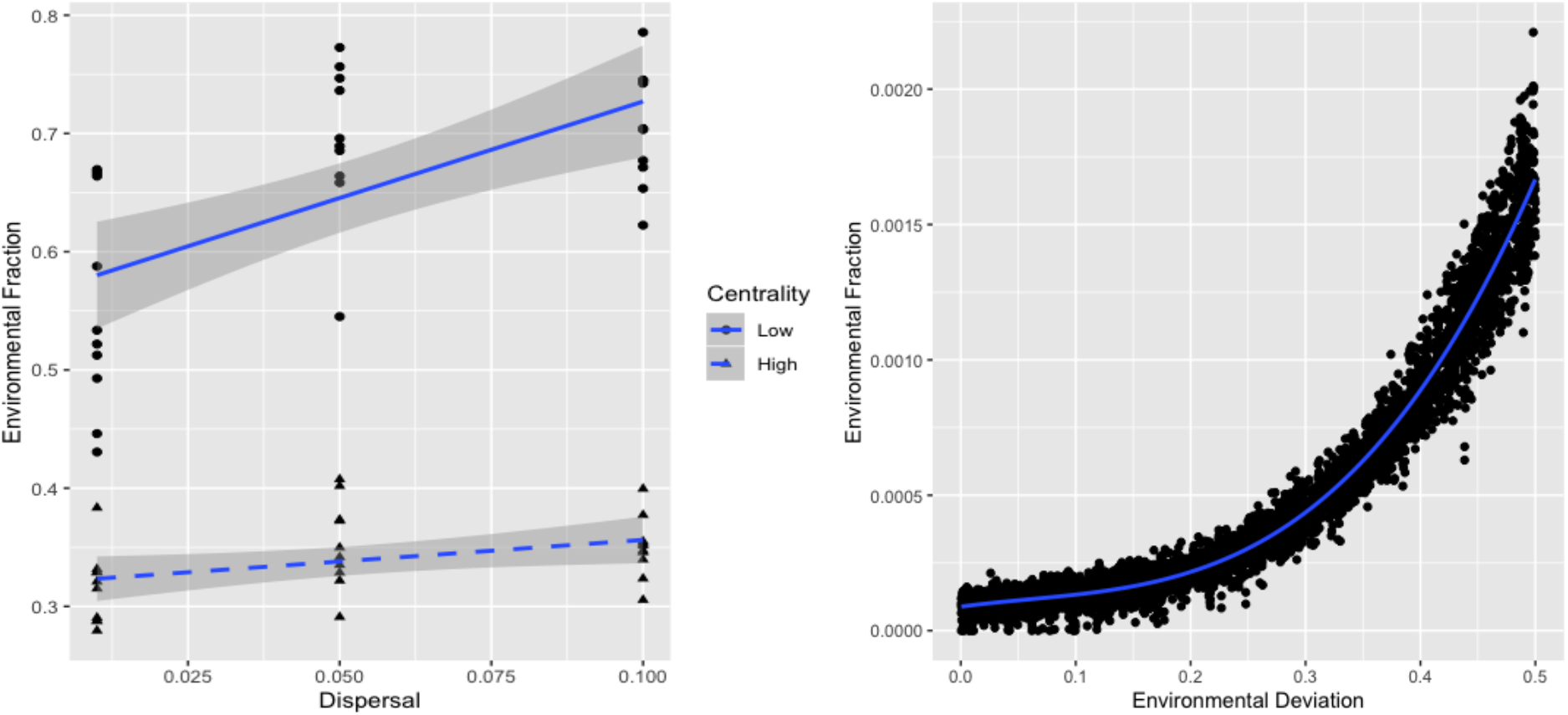
Effects of species traits (i.e. species optima; left panel) and site attributes (right panel) on the environmental fraction of variation in species distributions and site occupancy. A) Higher dispersal ability and lower niche centrality (i.e. greater deviation from mean niche value) enhance the degree to which different species (individual symbols) have distributions that correlate with environmental variation. B) Sites that differ more from the mean environmental value (environmental deviation) are more likely to be occupied by species with niche traits that are locally favored.

Sites also differed in how their species composition was related to environmental (fraction [E]) and spatial effects (fraction [S]) as well as co-distributions (fraction [C] *-* Figure 4b). Some sites tended to be occupied by locally dominant species (in the lower left of the ternary plot, nearer to fraction [E]), while others were occupied by species found in nearby sites (lower right of the ternary plot, nearer to fraction [S]). Some sites were also occupied by combinations of species that were differentially associated with each other regardless of environment or dispersal (upper apex of the ternary plot, nearer to fraction [C]). As can be seen in Figure 4b, there were also a wide range of intermediate conditions. A major driver of this variation are local environmental conditions, especially in relation to how distinct the local environment is from the overall mean environment of the metacommunity (Figure 5b).

We further investigated the structure of the species co-distribution (fraction [C]). This covariation can be directly attributable to species interactions because we explicitly model the processes underlying metacommunity dynamics. However, even in our model, species co-distribution may not directly link to pairwise interaction coefficients, but rather may emerge as a complex relationship between species interactions and environmental conditions (Cazelles et al. 2015, Blanchet et al. 2020). To illustrate this, we show the co-distribution among species as a heat map separately for each of the five individual simulations presented in Figure 4 and compared them to the actual interaction matrix that describes interspecific competition in our model (Figure 6; similar heat maps obtained with the other scenarios are shown in the Supplement Information). Despite the fact that the same interaction matrix was used for all five of these simulations, the resulting co-distribution patterns are inconsistent in their details. However, these matrices show that there is consistency in several features of the co-distribution. For example, they all share the predominance of strong negative correlations along the main diagonal that match the interaction matrix we used. They also share a strong ‘checkerboard’ pattern with alternating negative and positive co-distributions between species when these are ranked against their environmental optima. Given the simple scheme of species interactions we used (Fig. 5 and SI), these results are consistent with the predictions that direct interactions are stronger than indirect ones and tend to weaken with the number of links in indirect chains even if the details of these effects are less predictable (Cazelles et al. 2015).

**Figure 6:**
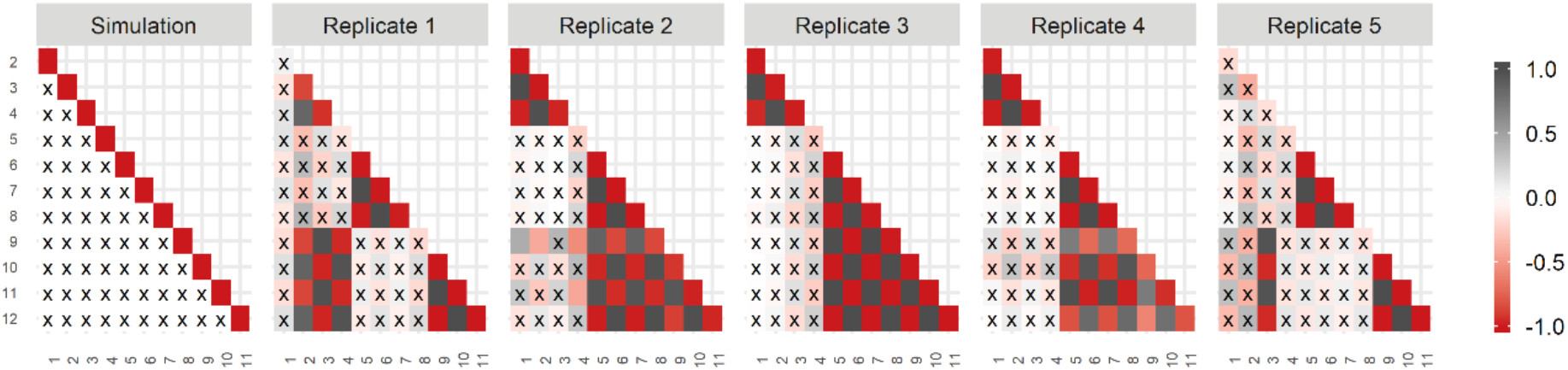
Comparisons of the interaction matrix (Simulation) with the co-distribution of species in five replicate runs (Iteration 1-5) of the scenario with interspecific variation in dispersal and competition among species. In each panel, species are ranked by the position of their environmental optima along the environmental gradient. The co-distributions are shown as heat maps with the strength of the covariation proportional to the intensity of color and the color indicating negative (green) or positive (gray) covariation among pairs of species. These can be compared to the pattern of direct species interactions (left panel called Simulations). The Xs denote no significant association although the color indicates the trend.

## Discussion

While metacommunity ecology has made great progress in the past decades, the assumptions that species and sites were relatively homogeneous in their underlying processes (reviewed in Leibold and Chase 2017), and the lack of explicit consideration of biotic interactions, have limited the applicability of metacommunity theory to relatively simple interpretations that have not addressed the dynamics of more realistic species pools within complex landscapes. Here, by combining a tractable process-based model with emerging analytical methods, we provide a general quantitative approach that accounts for multiple interacting assembly processes (including biotic interactions) that may operate differently among species or in different parts of landscapes.

Some of the issues we raise have already been highlighted in previous work that have shown that particular examples of species and site effects can occur, but here we sought to formulate a general analytical approach that accounts for these issues and test it against a process-based model. For example, Pandit et al. (2009) showed that species can be heterogeneous in their responses to environmental and stochastic factors depending on their degree of habitat specificity. Legendre and De Cáceres (2013) have calculated how sites and species contribute to beta-diversity (but not to the partitioning of driving factors). Others have argued that JSDMs can provide important insights into the drivers of such variation in species distributions (Hui et al. 2013, 2016, Pollock et al. 2014, Ovaskainen et al. 2017 see also Ovaskainen and Abrego 2020; but see Poggiato et al. 2021). Similarly, the heterogeneity among sites has long been identified as driving individual species distributions (see Guisan and Thulliers 2005, Soberon and Peterson 2005, Elith and Leathwick 2006) as well as driving overall variation among sites in global metrics of community structure (e.g. diversity patterns, etc.) as characterized by the field of landscape ecology (Turner 2005). Further suggestions that individual sites might vary in how they contribute to metacommunity dynamics include the concept of ‘keystone communities’ (Mouquet et al. 2012, Resetarits et al. 2017, Yang et al 2020) and metacommunity approaches to spatial networks (Economo and Keitt 2010, Borthagaray et al. 2015, Harvey et al. 2020). These disparate approaches (species vs sites) to metacommunities are doubtlessly closely related to each other and can be linked by the emerging methodologies of methods such as JSDMs to develop a more nuanced metacommunity ecology that recognizes a plurality of mechanisms and processes underlying community assembly. The expectation here is that the internal structure framework can provide further insights on these complexities.

Although we find that JSDMs can reveal important aspects of the internal structure of metacommunities, we find that there are some remaining important challenges to resolve in quantitatively making process-pattern linkages in metacommunities (see also Poggiato et al. 2021, Miele et al. 2021).

Although there are some important challenges to consider, our study illustrates important insights about the internal structure of metacommunities, including:

1. Variation partitioning using JSDMs (here implemented using HMSC) can be a useful tool to describe how basic processes of community assembly at the species and sites levels (e.g. environmental selection, dispersal, biotic selection, and drift) might determine metacommunity wide variation in community composition and thus link the internal structure of the metacommunity where these processes act to the external structure that summarizes these effects at the broader spatial scale.
2. Quantifying co-distributions of species in metacommunities can improve predictive ability even when the processes that generate these distributions are complex (stochasticity, complex spatial landscapes, and species interactions).
3. Species can have distributions that vary in the degree to which they are determined by combinations of the basic community assembly processes depending on features of their ecology (e.g. dispersal and environmental preferences); and
4. The predominant assembly processes that determine local communities can differ among adjacent sites in a metacommunity (e.g. sites that are occupied by species most fit for environmental conditions vs sites occupied by species that do well in nearby sites due to dispersal even though they differ in environmental conditions).

It is important to emphasize that there remain some substantial challenges in moving forward with the overall approach we advocate in this paper and producing more robust versions of the internal structure framework for metacommunities. These include technical issues, such as the estimation of parameters and interpretation of results in more complex models (e.g., the interaction between dispersal and environment such that species are limited in their dispersal abilities due to environmental features of the landscape matrix), as well as conceptual ones, such as accounting for other processes such as speciation, local adaptation, and historical biogeography. Nevertheless, we see that producing robust frameworks taking into account the internal structure of metacommunities will be fruitful, allowing a deeper understanding of ecological dynamics in more realistic, but necessarily complex, spatial landscapes.

Our analytical framework simplifies several potentially complex processes (e.g. non-linearities and interactive effects of mechanisms) into an approximation involving colonization-extinction dynamics. It is possible that more realistic and complex mechanisms driving these processes will weaken associations between pattern and process or create biases in the partitioning of the variation revealed by JSDMs. However, the developments of JSDMs are still progressing, and we anticipate that future developments will solve some of these problems (see Wilkinson et al. 2020).

The co-distribution component of the JSDMs (fraction [C]) is particularly concerning. We find that the component estimated by the JSDM often deviates from the settings of the process-based model, especially when species have broad environmental niches (Figure 2). A possible reason could be sensitivity that leads to biases in the HCSC fitting procedure, but the more likely explanation is that the [C] fraction, additional to true biotic interactions, tends to absorb any process that is inadequately quantified by the environmental (fraction [E]) and spatial components (fraction [S]) (see Blanchet et al. 2020). In addition to species interactions, this would include, for example, unmeasured environmental factors (see Blanchet et al. 2020), inadequately quantified landscape attributes, or that the process-based model creates somewhat different environmental responses than assumed in the JSDM. Teasing apart the effects of species interactions from these confounding factors should thus be a major focus for future work. Nevertheless, it is important to understand that including the co-distribution component in our analyses allows us to account for them, rather than lumping them with residual variation where they have likely given a greatly exaggerated impression of stochasticity.

Our framework links most naturally to mechanisms that focus on interspecific competition, in analogy with evolutionary genetics (Vellend 2010, 2016). However, species interactions in metacommunities are much more variable and include consumer-resource interactions, mutualisms, and facilitative interactions. Although such interactions can easily be incorporated in simulations, the interpretation that might link process to pattern in such cases are likely to become more complex (see Gravel and Massol, 2020). Likewise, future work could include local (co-)evolutionary dynamics (see Urban et al. 2020) and historical effects of biogeography and speciation (Leibold and Chase 2018, Overcast et al. 2020). Here, we have also retained a simple two-level perspective on spatial scale (local discrete sites in a broader regional landscape). It is increasingly apparent that metacommunity dynamics occur over multiple nested scales and that habitats can be continuous and/or nested, rather than discretely patchy (e.g. Munkenmuller et al. 2012, Rybicki et al 2018, Ovaskainen et al. 2019, Viana and Chase 2019, König et al. 2021). Refining our approach to address multiple spatial scales is a logical next step.

Finally, it is increasingly clear that temporal dynamics of community change in metacommunities can provide critical insights about the mechanisms that drive metacommunity patterns (e.g. Jabot et al. 2020, Blanchard et al. 2020, Guzman et al. 2021). We imagine future work on the internal structure of metacommunities as being very amenable to incorporating temporal changes (see for example, Ovaiskainen et al. 2017 for an initial step in this direction). Here we have focused on purely spatial approaches because there are still remarkably few studies that might permit sufficiently structured data to permit temporal analyses and because the limitations and challenges of such analyses are not yet clear.

In conclusion, we have argued focusing on the internal structure of metacommunities, by examining site-specific and species-specific variation components, and better accounting for biotic interactions in community assembly can enhance our ability to infer process from pattern in the distributions of species and the occupancies of sites. We also argue that continued work on this focus is essential if metacommunity ecology is to address the dynamics and structure of realistic metacommunities that have typically high biodiversity and occur in complex landscapes. This focus on internal structure represents a shift from traditional approaches that used descriptors of overall variation components at the metacommunity scale that generalizes previous ad hoc approaches to similar internal variation in metacommunity patterns (e.g. Pandit et al. 2009, Legendre and De Cáceres 2013). The dynamics and structure of distributions of realistically diverse species in a realistically structured landscape of sites likely involves the interaction of community assembly processes including environmental filtering, dispersal, and drift, and these are unlikely to be adequately described by simple metacommunity level metrics (see Ovaskainen et al. 2019). Consequently, dissecting the internal structure of metacommunities on the basis of species and site contributions could provide key insights into the processes underlying metacommunity assembly. Such insights might be particularly useful in the management of landscapes and metacommunities for conservation purposes since they focus on particular units (species or sites) that are often the focus of concern in such cases.

### Speculations

Our goal in this paper was to emphasize that looking at species and site specific differences (what we call the internal metacommunity structure) may reveal more nuanced information about the underlying processes than using the global summary statistics that have been the focus of most of the previous work. Using JSDMs is one way in which estimates of this internal structure can be generated, and they have the additional advantage that they estimate species associations, which can be related to biotic interactions. We suspect, however, that other approaches can be developed that might be equally (or possibly even more) useful than JSDMs for the general purpose of this paper, especially if they would be able to include also dynamic mechanisms such as dormancy (Wisnoski et al. 2019), local evolution of ecological traits (Urban et al. 2008), and ecosystem-level feedbacks (Loreau et al. 2003). The challenge to consider such processes is to include them into an statistical framework that is still comparable in simplicity and general applicability to the JSDM framework that we used here. It is worth noting that any method could be adapted with different degrees of difficulties to the framework of internal structure of metacommunities.

## Acknowledgements

This paper emerged from workshops funded with the support (to JMC) from the German Centre for Integrative Biodiversity Research (iDiv) Halle-Jena-Leipzig funded by the German Research Foundation (FZT 118). MAL would like to acknowledge the Alexander von Humboldt Foundation for a Research Award and NSF award 2025118 that helped fund this work. LDM acknowledges KU Leuven Research Fund project C/2017/002 and FWO projects G0B9818 and G0C3818. We also thanks F. Engel, C. Rakowski, K. Taylor, R. Pelinson, X. Zhao, L. Juen, T. Michelan, A. Rudolf and D. Jenkins as well as an anonymous reviewer for comments on earlier versions of the manuscript.

## Supplementary Information

### Description of Model

#### Patch locations and environmental variables

In our model, the metacommunity consists of *N* patches distributed over a spatially heterogeneous landscape, with multiple environmental variables (although, the current simulations only have one environmental variable *D* across 1000 patches) that could either be randomly distributed or spatially autocorrelated. Each patch has a set of coordinates in a two-dimensional space, and all possible coordinates are feasible such that this is a continuous space model that is not restricted to a lattice or some other kind of regular spatial arrangement of spatial units. A patch may be empty or be occupied by a single or by several species. We define *X*_*i,z,t*_ as a stochastic variable representing the occurrence of species *i* at location *z* and time *t*. Occurrence, *X*_*i,z,t*_, takes a value of 1 when species *i* is present and a value of 0 when it is absent. Similarly, we define *Y*_*z,t*_ = *X*_1,*z,t*_, *X*_.,*z,t*_, …, *X*_*R,z,t*_ as a vector containing the presence-absence of each species from the regional pool *R*.

The model only tracks patch occupancy (not population densities). Spatial dynamics occurs because of colonization events, in both empty patches and patches that are occupied by other species, and because of extinction events. The emerging species co-distributions are a result of a dynamic balance between these events. Ecological interactions can impact either or both the colonization and the extinction probabilities. For instance, the presence of a competitor pre-empting a patch can reduce the colonization probability by another competitor. Alternatively, the presence of a competitor in a patch could increase the extinction probability of another species. Similarly, the environment could influence both the colonization and the extinction probabilities.

#### Patch Colonization

We consider a discrete-time Markovian process to represent the dynamics of presence-absence of all species and we incorporate the effect of dispersal, environmental filtering and ecological interactions in such a way that we could cover all possible scenarios wherein species differ in any combination of these mechanisms and processes. We can include interspecific competition along with other types of spatial dynamics such as predator-prey interactions (Gravel et al. 2011), priority effects (Shurin et al. 2004), or mutualistic interactions (e.g. Gilarranz et al. 2015). In this paper, we focused on competition only though. Following a colonization event from time *t* to *t* + *Δ* corresponds to:

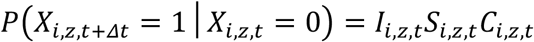

where *I*_*i,z,t*_ is the number of immigrants of species *i* reaching patch *z* at time *t, S*_*i,z,t*_ is the effect of environmental filtering on the probability of establishing a viable local population and *C*_*i,z,t*_ is the effect of ecological interactions on the establishment probability. We note that because we represent a stochastic process, the product of these three functions has to be bounded between 0 and 1. We consequently define these quantities:

The effect of immigration is given by:

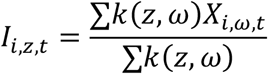

which is a weighted average of the occurrence probability of species *i* in the neighborhood of *z*. The function *k*(*z, ω*) is a dispersal kernel that depends on the location of patch *z* and the neighborhood *ω*. For convenience, we considered an exponential function of the Euclidean distance between localities. We added to the kernel a low distance and neighborhood-independent constant *m* to account from immigration from outside the simulated metacommunity. This assumption is required to prevent total extinction by drift under pure neutral dynamics.

The effect of the environment is given by a product of the establishment performance over all environmental variables *E*_*n*_:

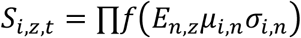

In our simulations, for convenience, we considered that the function *f* has a quadratic form for all species and all environmental variables, though the model is flexible and general enough to consider other (non-linear) responses that could also differ among species.

#### Ecological interactions on establishment probability

To incorporate all possible ecological interactions, we started by representing the interaction network by a community matrix *A* of *R* species. The elements *α*_*ij*_ of *A* quantify the effect of species *j* on the dynamics of species *i*. When *α*_*ij*_ is negative, the colonization probability of species *i* decreases and/or its extinction probability increases when *j* is found locally. Inversely, when *α*_*ij*_ is positive, the colonization probability increases and/or the extinction probability decreases. To account for the cumulative effects of local interactions on transition probabilities, we made colonization and extinction probabilities community dependent. As explained above, at a time *t*, the *Y*_*z,t*_ vector gives the local assemblages. We calculated the sum of interactions at any time and for each species as *ν* = *A*_*z,t*_*Y*_*z,t*_. Our approach can be interpreted as a spatial analogue to the generalized Lotka–Volterra model because it takes into account the impact of the whole network of interactions on each species dynamics and can deal with any type of interaction. We now define the function:

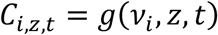

representing the total effect of ecological interactions on the colonization probability. For convenience, we will use a sigmoid function, with *g* ranging between *c*_*min*_ at high negative interactions and *C*_*max*_ at high positive interactions, where *c*_*max*_ should be interpreted as the maximal colonization probability when the environmental conditions are optimal and there are no dispersal limitations.

#### Patch Extinction

The definition of the extinction probability follows exactly the same rules as for colonization, except that extinction is independent of the neighborhood composition. We follow the same logic to define the effect of ecological interactions and of variation in the environment. Consequently, we get the Markovian process:

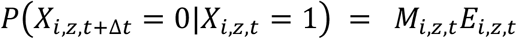

where *M*_*i,z,t*_ and *E*_*i,z,t*_ are the responses of the extinction probability to the local environment and to ecological interactions, respectively. The difference with the colonization functions defined in the previous section is that the extinction probability must be larger when interactions are negative and smaller when they are positive. In addition, the extinction rate should be minimal (instead of maximal) at environmental optimum.

#### Interpretation

To interpret the model, note that, at steady state, for each species, we obtain the expected occurrence probability (*P*^) at each site as:

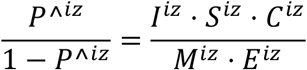

After a log transformation, this yields:

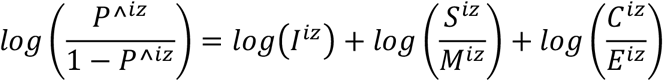

This last equation can be interpreted as a macroscopic description of the expected species distributions pattern (Thuiller et al. 2013). In this formulation, *log*(*I*) describes the tendency of a patch to resemble other nearby patches due to the spatial contagion by dispersal, 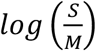 describes the tendency of sites to be occupied by species with similar fitness responses to environmental gradients, and 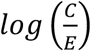 describes the remaining influence of other species on co-occurrence due to interactions among species. The values for these indices will depend on what choices are made for the components of eq. 1 (see Supporting Information for details on how we implemented this simulations model).

This modeling framework can represent the classical archetypes but also permits more intricate (and likely far more realistic) metacommunity scenarios and predictions. For example, we could use the model to examine how species traits (and environmental context) link to metacommunity dynamics. Moreover, continuous mixtures of different metacommunity extremes (archetypes) can be represented by appropriate parameter choices for dispersal, competitive abilities, and environmental preferences. For instance, species sorting would require a relatively large colonization to extinction ratio along with species-specific environmental requirements and regional similarity (sensu Mouquet and Loreau, 2002). Alternatively, coexistence within competition-colonization trade-offs requires species to have similar responses to the environment and appropriate heterogeneities in the *I, C* and *E* functions, but no environmental preferences.

The implemented mechanisms in the simulation model can be partially mapped onto variation partitioning components. For instance, at equilibrium, we could expect dispersal limitation (the *log*(*I*) term in equation 3) to create positive spatial autocorrelation at the dispersal scale (the [S/E] fraction in variation partitioning, i.e., spatial variation independent of environmental selection). Environmental selection (the 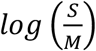 term in the last equation) should lead to a correlation between composition and environment (the [E/S] fraction in variation partitioning). The last term in equation 3, however, describing the effect of interactions on distribution (the 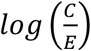), is novel and has no equivalent in the context of classical variation partitioning.

There are some interesting properties to point out regarding our proposed variation partitioning scheme. First, by considering the combined effects of environmental selection, dispersal and interactions, the final residuals (unexplained sources of variation) in the model leading to this new partition variation scheme is (in principle) solely related to non-spatialized independent species variation. Second, in our variation partitioning, the interaction component is due to species co-variation (i.e., a joint component among species distributions). In empirical community data, however, this interpretation can only be made if all the environmental variation (predictors) underlying environmental selection in empirical community data has been incorporated (as pointed out in the main manuscript). If not, then the spatial and species interaction components could be measuring variation related to unmeasured environmental variables that are either spatialized (i.e. characterized by the spatial component in variation partitioning) or shared among species (i.e. joint component).

### Description of the Statistical Framework

#### Hierarchical Community Models

In their simplest form, Hierarchical Community Models (HCMs) resemble standard species distribution models that regress species presences/absences against environmental predictors (i.e., logit link). However, to reduce model complexity, HCMs assume that all species in a metacommunity will react to environmental heterogeneity following a similar response function (e.g., linear vs quadratic or Gaussian). The same assumption is made in common variation partitioning (see Peres-Neto et al. 2006). To model the spatial component (i.e., due to spatialized dispersal), either spatial variables such as Moran’s eigenvectors maps (MEM, Dray et al. 2006) or spatially auto-correlated latent variables (Ovaskainen et al. 2016b) can be incorporated to the model. To account for biotic interactions, non-spatially auto-correlated latent variables are used. If we use a linear specification approach (here, this can also include quadratic terms that capture Gaussian responses to environment as imposed in our model), we can write:

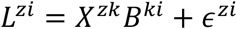

with

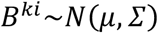

where *L*^*zi*^ is the presence (or absence) of species *i* (out of *m* species) at patch *z* (out of *n* patches), *X*^*zk*^ is the value of the environmental variable *k* (out of *p* variables) at site *z, B* is a matrix of regression parameters, *µ* is a vector of length *p* that describes the mean environmental response of all species, *Σ* is a *p* × *p* covariance matrix that describes how species vary (diagonal) and co-vary (off-diagonal) around the mean environmental response (Ovaskainen and Soininen 2011), and *ϵ*^*zi*^ is a residual value. Estimating species parameters hierarchically around a community mean reduces the degrees of freedom and makes the model easier to fit with limited data. Note that both *µ* and *Σ* can be further informed or constrained by species traits or phylogeny if desired (Ovaskainen et al. 2017). To account for biotic interactions, we consider latent variables *H*^*zl*^ (where *l* refers to a latent variable measured at site *z*) and their associated parameters *Λ*^*li*^ (Ovaskainen et al. 2016a). This yields:

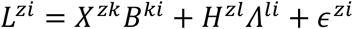

Note that it is not necessary to always include all of these components in one model; they can be considered in any combination deemed relevant for a particular question. In this paper, we used Moran’s Eigenvector Maps (MEMs; Dray et al. 2006), a powerful and commonly used method to model spatial autocorrelation in statistical models involving species distributions.

#### Calculating Variation Partitioning for the HCM

As in any generalized linear mixed effect model, we can now partition the explained variation into different components, notably environmental heterogeneity, space, co-distribution (biotic interactions), and unexplained variation (Figure S1). To estimate the contributions of each of these four fractions for each species, we calculated semi-partial coefficients of determination (i.e., based on Type III sum-of-squares as specified in Peres-Neto et al. 2006) using the implementation suggested by Tjur (2009) as being more appropriate for presence-absence data (i.e., logit link) than the traditional variation partitioning based on an identity link. To adjust for the number of variables used to quantify each fraction of the variation partitioning analysis, we applied the adjustment to the coefficient of determination proposed by Gelman and Pardoe (2006) in the variation partitioning analysis, which is designed for hierarchical models. As shown in Figure S1, the different fractions were combined so that a unique value was associated to environment (fractions [a], [d]/2, [f] and [g]/2), co-distribution (fraction [c]), space (fractions [b], [d]/2, [e] and [g]/2) and the unexplained portion of the variation (fraction [h]). Latent variables are quite powerful to isolate structure in the data. As such, in the calculation of the variation partitioning, latent variables will capture almost all (if not all) variation associated to the environment and space, giving an artificial inflation of the overlapping partitions between co-distribution and environment and co-distribution and space. For this reason, all partitions overlapping with co-distribution (fractions [e], [f] and [g]) were assigned to either environment (fractions [f] and [g]) or space (factions [e] and [g]). In this calculation, a unique measure of explained variation (akin to adjusted R^2^) is associated to co-distribution (fraction [c]) but this is not the case for environment and space. To associate a unique value to environment and space, and represent the results as we did in Figure 2 and 3 (main manuscript), we divided the fractions overlapping environment and space between these two components. As such, the sum of fractions [a], [f], half of fraction [d] and half of fraction [g] were used to measure the effect of the environment while fractions was considered [b], [e], half of fraction [d] and half of fraction [g] were used to measure the effect of space. This scheme in which half of common variation is assigned to two or more common components is commonly used in hierarchical partitioning (Chevan & Sutherland 1991).

**Supplementary Figure 1.**
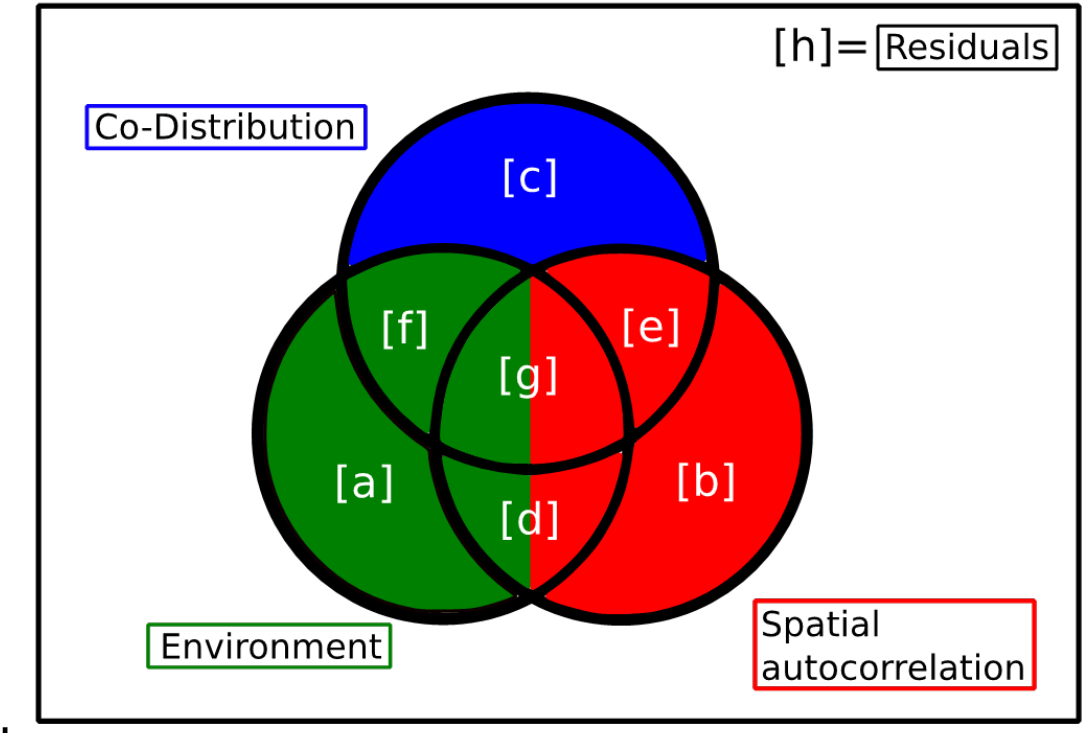
variation partitioning scheme used to estimate the importance of each matrix of predictors.

#### Calculation of the coefficient of determination

##### 1) Classic coefficient of determination

The coefficient of determination, *R*^2^, that was partitioned in the variation partitioning analysis (Appendix XX) is calculated for any given species *j* as:

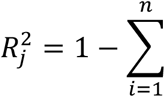

where *y*_*ij*_ is the data (presence-absence) associated with species *j* (out of *p* species) at site *i* (out of *n* sites), *y*^_*ij*_ is the model (predicted value) associated to species *j* at site *i* and <u>*y*</u>_*j*_ is the average of the data (i.e., sum of presences divided by *n*) for species *j* across all sites.

##### 2) Community-level coefficient of determination

Although having an 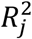 for each species *j* can be highly informative and is part of our framework on the internal structure of metacommunities, it can be also useful to estimate the contribution of single communities *R*^2^ to the entire metacommunity. This is obtained by averaging all 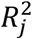:

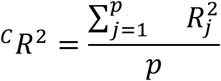

where the,^*C*^*R*^2^ is the community-level *R*^2^.

##### 3) Site contribution to the coefficient of determination

In the paper, we use the contribution of each site to 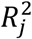 to present how each site contributes differently to the environment, space and co-distribution for the community. The calculation of the site *i* contribution to the 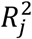, is calculated as:

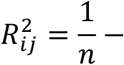

The first part of the equation where the 1 of the classic 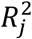 is divided by *n* is included to make sure that if we sum all 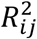 across all sites for species *j*, the resulting value equals to 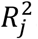.

More importantly, what can be noticed is that by calculating 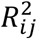, the contribution of sites to each species 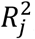, we obtain a matrix that has the same dimension as the site by species matrix (an *n* × *p* matrix). Using this matrix, if we sum across all sites, we obtain the 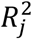.

However, if we average across the species for site *i* we obtain the site’s contribution to the community-level ^*C*^*R*^2^ or 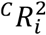.

The amount of variation expressed by the 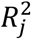, the ^*C*^*R*^2^, the 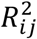 or the 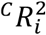 can all be partitioned in its environmental, spatial and co-distribution component following the procedure presented in the section “Calculating Variation Partitioning for the HCM” above.

### Parameterization and Simulation Scenarios

We simulated metacommunity dynamics with a landscape of 1000 patches over 200 time steps and an initial occupancy of 0.8. Previous studies show that steady state dynamics are obtained substantially earlier than 200 time steps with this initialization and we use this to interpret our results. Patches were placed randomly in a two-dimensional plane with coordinates drawn from a uniform distribution with a minimum of 0 and a maximum of 1. The environment varied spatially, with values drawn from a random distribution between 0 and 1. In the specific simulations we studied in the paper, colonization was the only component of the species that were affected by the environment (i.e. *E*_*i,z,t*_ = 1). Specifically, colonization reacted to the environment following a quadratic curve.

For all scenarios considered, we simulated 12 species. Niche optimums for the species were evenly distributed between 0.1 and 0.9 while niche breadth was set to 0.8 for simulations with narrow niches (scenarios A, B, E, F, G), and to 2 for simulations where niche was assumed to be broad (scenarios C and D). For dispersal, we considered an exponential dispersal kernel, with a distance-independent immigration probability of 0.001 and an *α* parameter of 0.05. For scenario G, where we have variable dispersal kernels (Figure 3), *α* was 0.01 for 1/3 of species, 0.05 for 1/3 of species, and 0.1 for the other 1/3 of species. We used a sigmoid function to relate the total number of interactions with colonization and extinction coefficients following the implementation by Cazelles et al. (2016). Colonization probability in the absence of interactions was set at 0.4, which tends to zero as negative interactions tend to infinity, while it asymptotes at a 1 with infinite positive interactions.

All other aspects of the colonization-interaction curve were the same for all scenarios. Similarly, extinction in the absence of interactions was set at 0.025, and tended to 1 with infinite negative interactions, while its asymptote tended to 0 with infinite positive interactions. In both cases, the parameter setting the shape of the sigmoid function was set to 0 for the scenarios without competition (scenarios A, B, and F), and 1.5 in the presence of competition (scenarios C, D, E, G) . If there were interactions, then a focal species only interacted with the two species that had the closest niches. For all scenarios, five sets of metacommunities were simulated and analyzed to obtain the results found in Figures 2, 3, and 4 in the main text, and supplementary figures.

All scenarios have been implemented in R and the project’s repository can be found here: github.com/javirudolph/testingHMSC

**Supplementary Figure 2.**
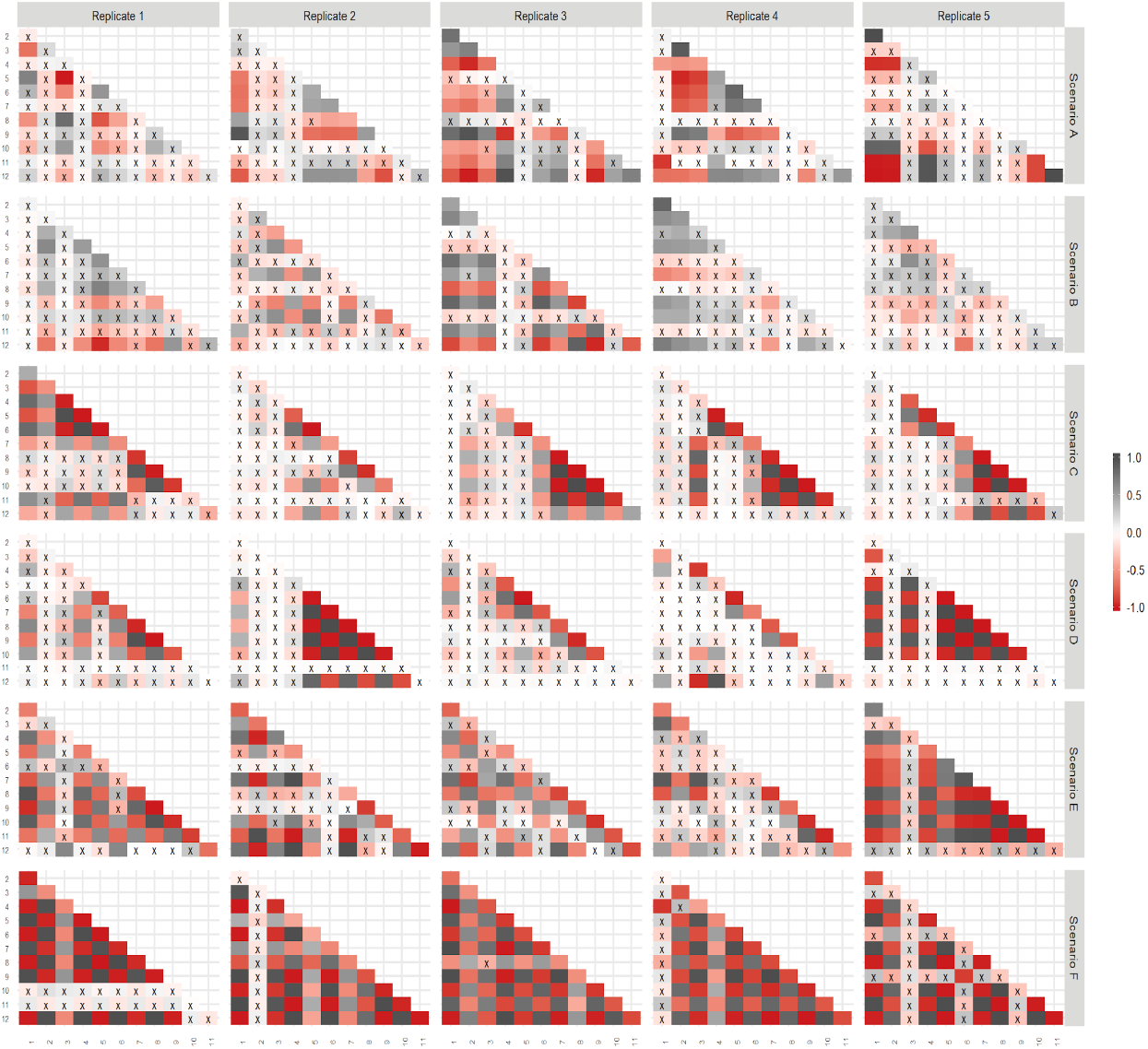
Species interactions for all scenarios.

**Supplementary Figure 3.**
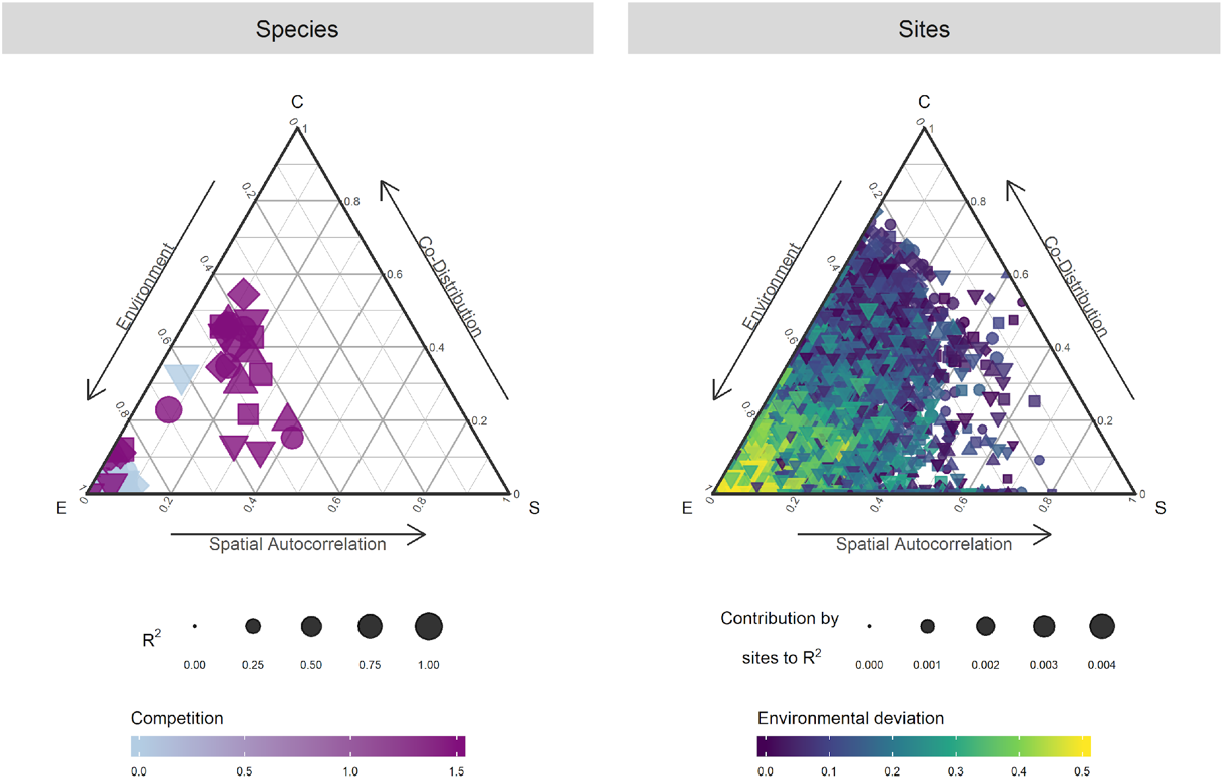
In which we have half of the species with interactions and the other without.

**Supplementary Figure 4.**
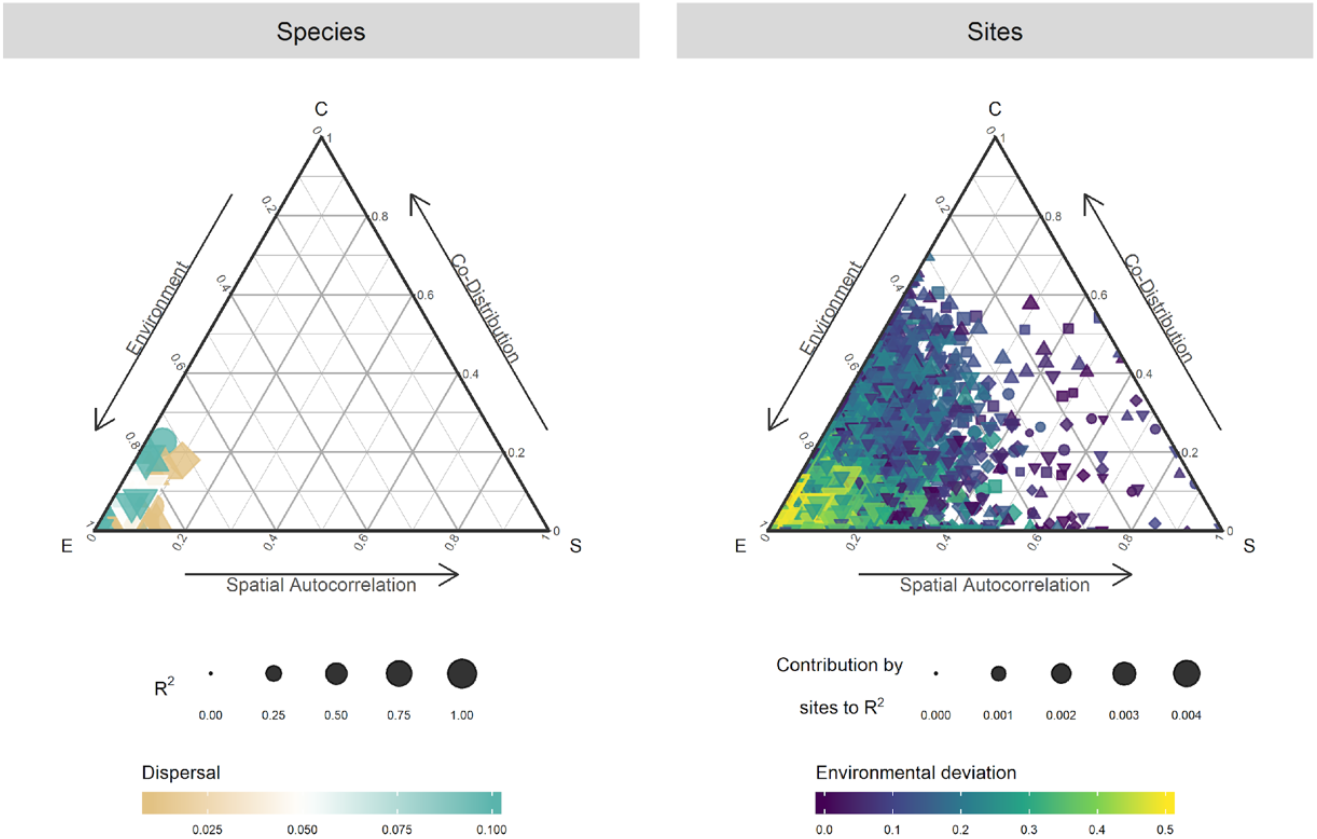
In which we change dispersal only, *α* was 0.01 for 1/3 of species, 0.05 for 1/3 of species, and 0.1 for the other 1/3 of species.

### Example of metacommunity simulation functions and analyses

The following example of code shows the overall processes involved in the metacommunity simulation for our model. This example shows the scenario for 20 patches and one environmental variable. The model gives the option for a random or spatially aggregated structure for the patches. In the aggregated case, we determined four clusters, denoted by `Nclusters` in the code below. The value of the environmental variable for each patch is shown with the color hue. In this case, the environmental variable is randomly distributed.

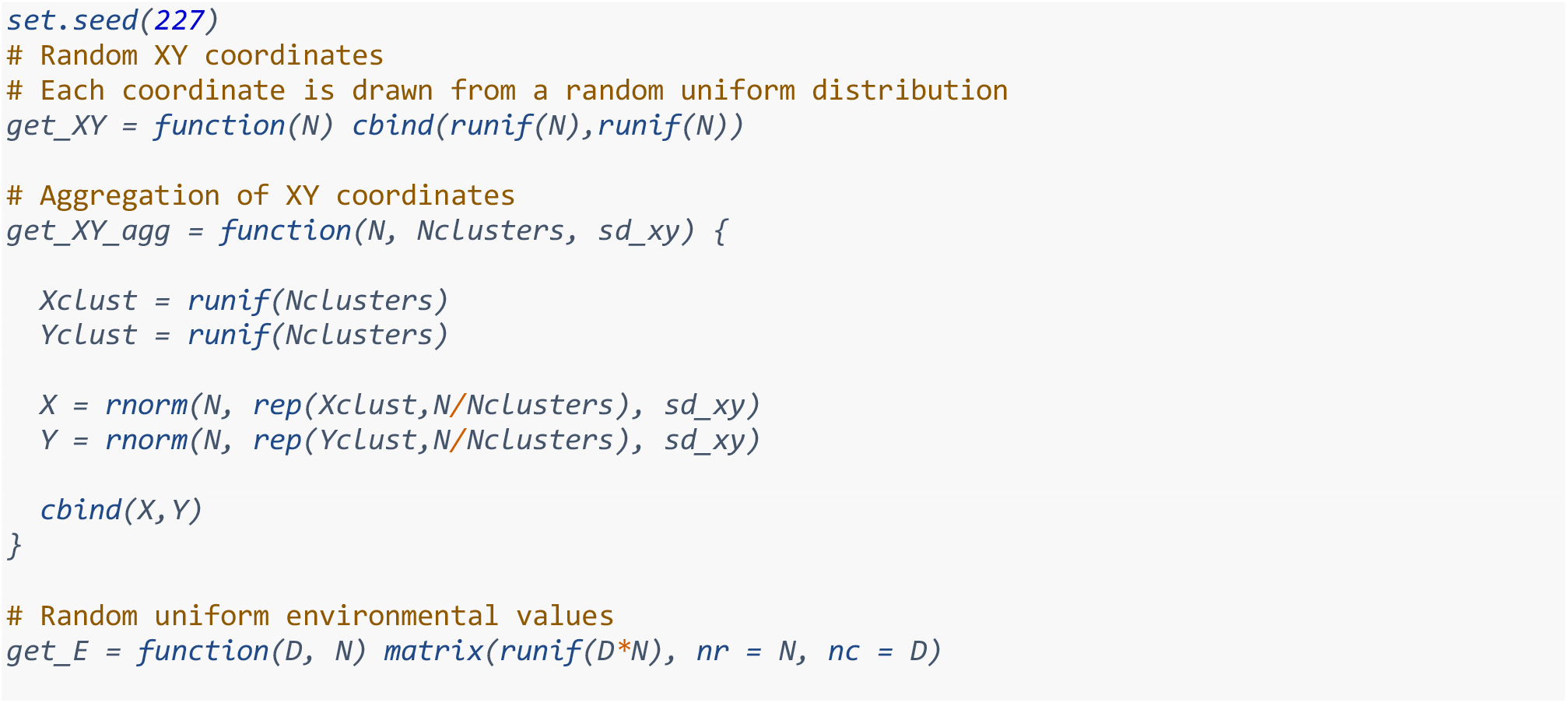

Here, we set N, the number of patches to 20, and D, the number of environmental variables to one.

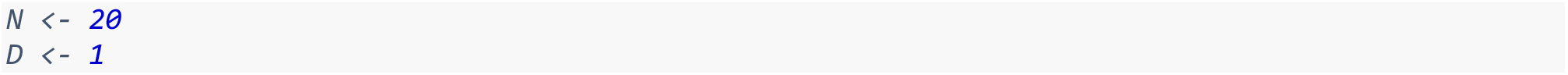

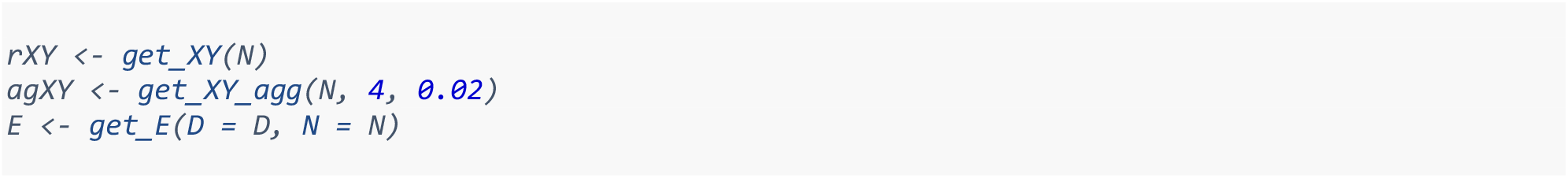

The following figure shows a side by side comparisson between random and aggregated patches obtained from our functions.

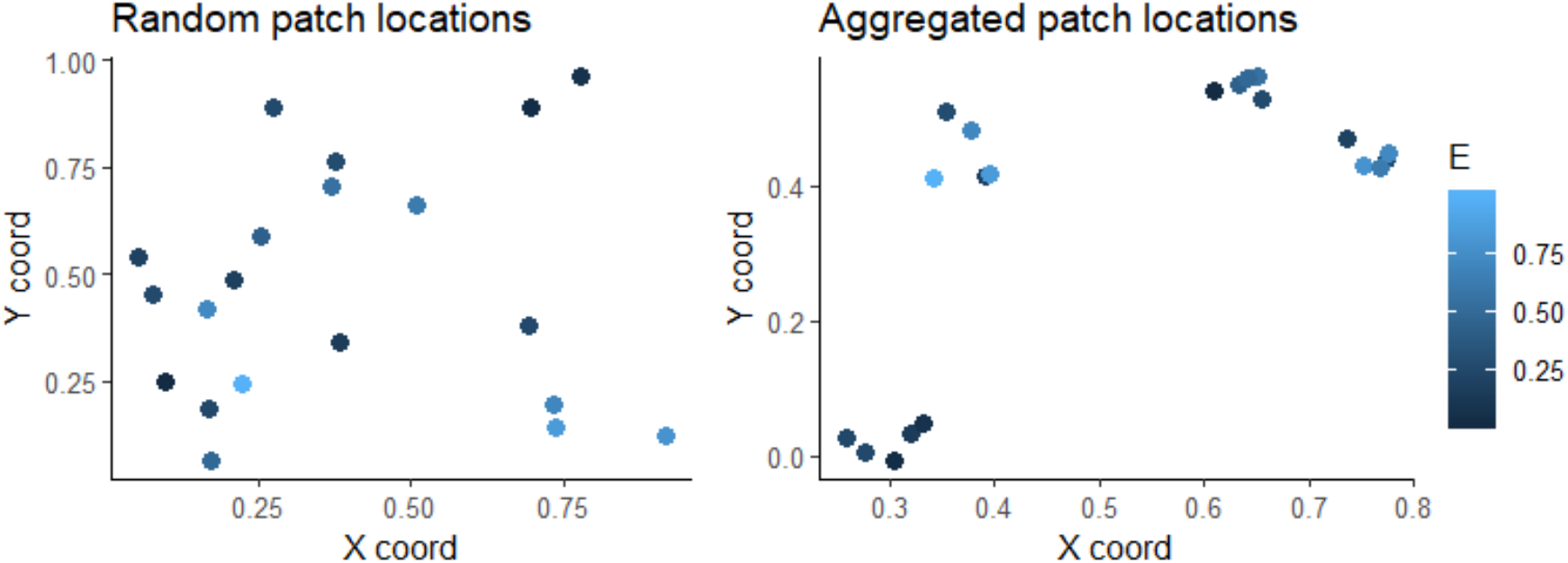

#### Initial Occupancy

The toy model allows for setting the initial occupancy in the metacommunity. For example, to create the initial conditions, *t* = 0, of presence absence, species occupancy is drawn from a random uniform distribution, and values smaller than 0.8 are considered as species presence. Patches or locations *z* are represented by rows in the matrix, whereas each species is a column. Each cell in the matrix is *X*_*i,z,t*_ and each row is *Y*_*z,t*_ for *t* = 0. The following figure shows three different iterations of this process, where areas in black represent occupancy = 1, and white denotes an absence.

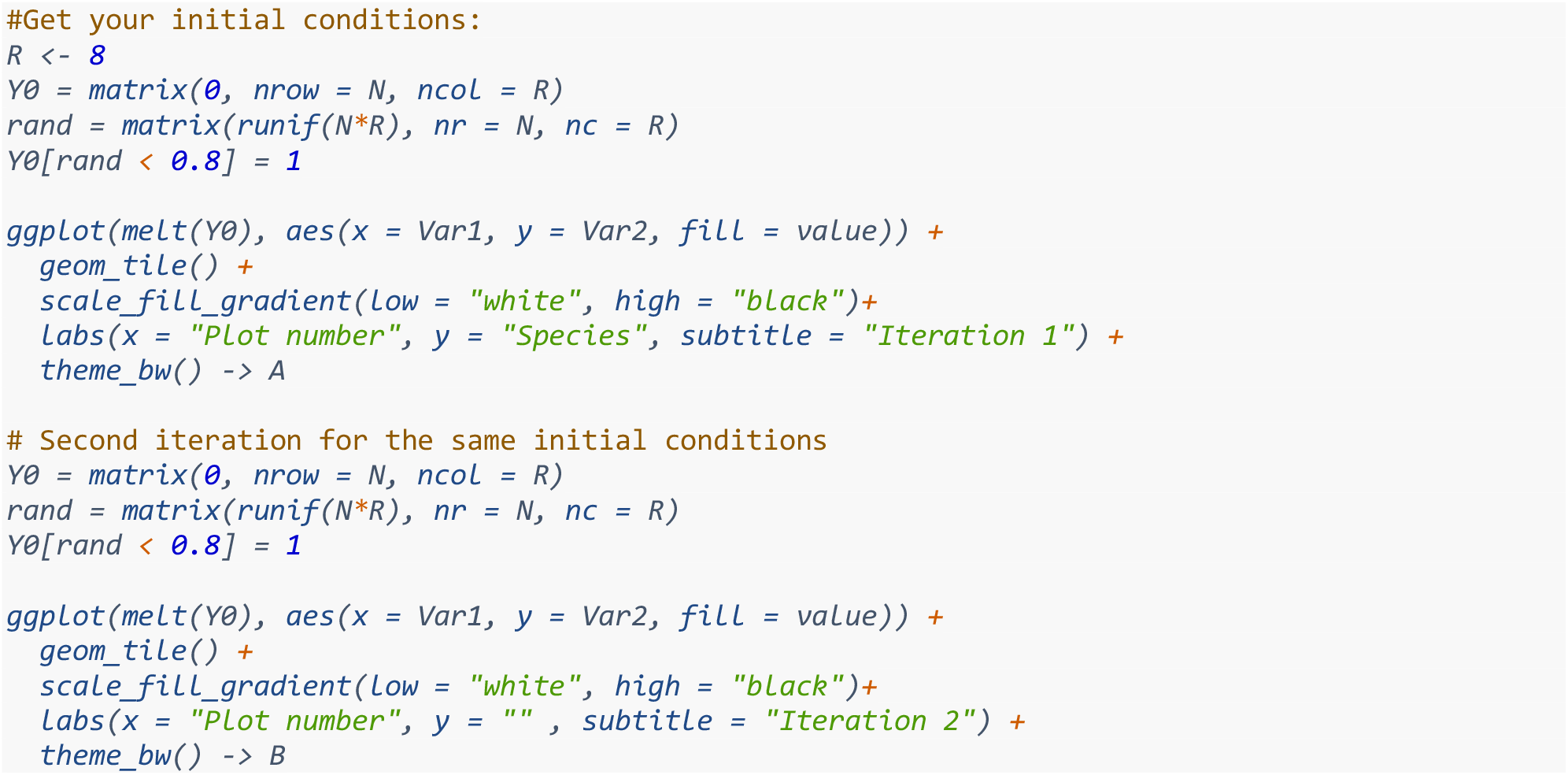

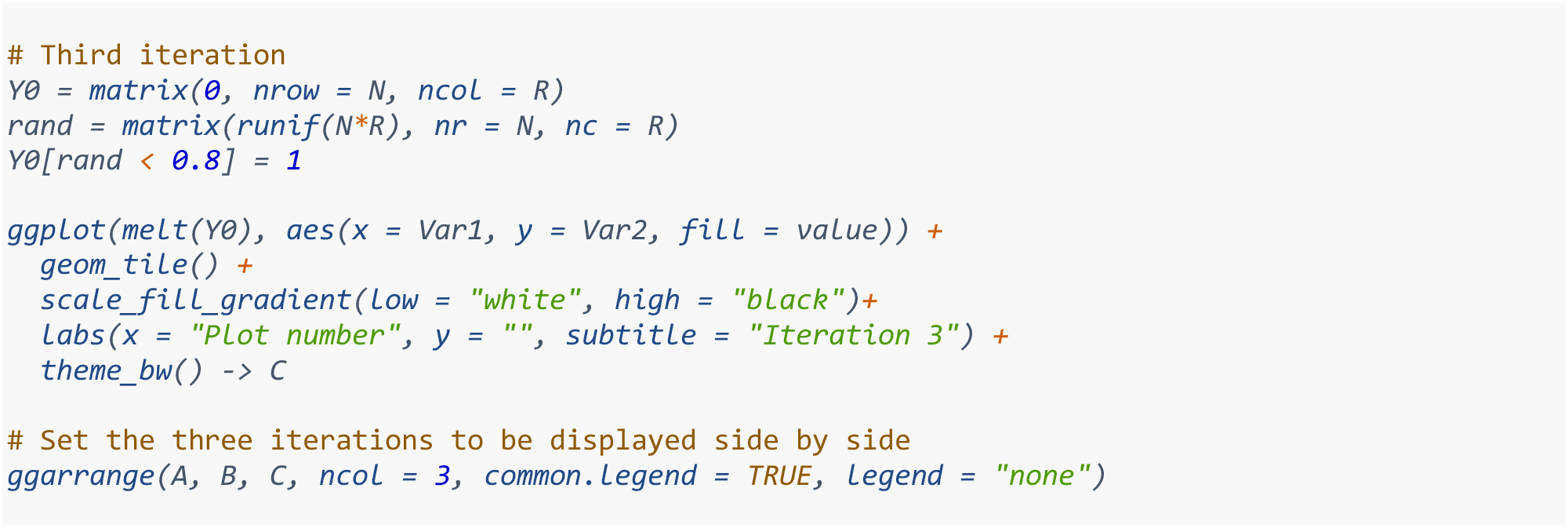

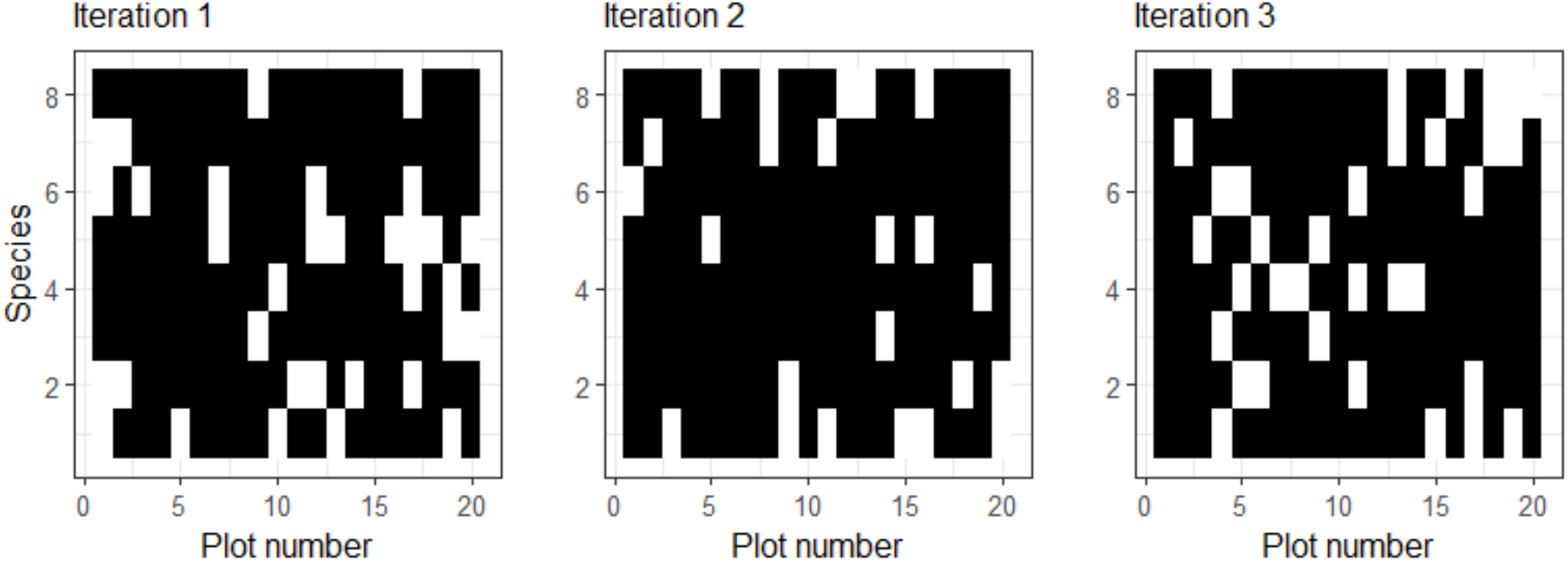

When we consider the immigration component, we need to also consider the connectivity matrix. In the code below, `I_f` calculates the probabilty of immigration for each species, based on the occupancy matrix and the dispersal kernel, K. The argument `K` is the connectivity matrix. The argument `XY` corresponds to the patch coordinates, whereas `alpha` is the dispersal parameter associated to the exponential distribution used for dispersal. It can be computed with the following function:

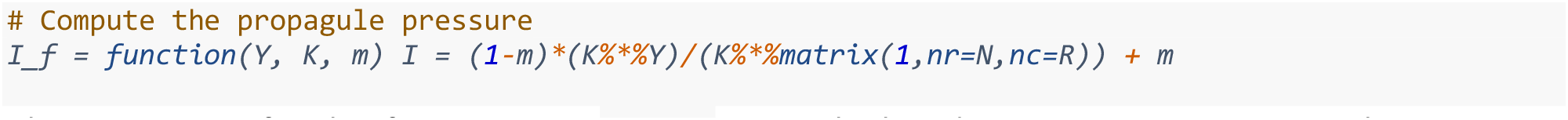

The arguments for this function are: `Y, K, m`. We calculated *Y*, species presence or absence, in the previous section with the case for initial conditions. Argument `m` is set in the parameters as a value *m = 0*.*001* and the connectivity matrix `K` is calculated below.

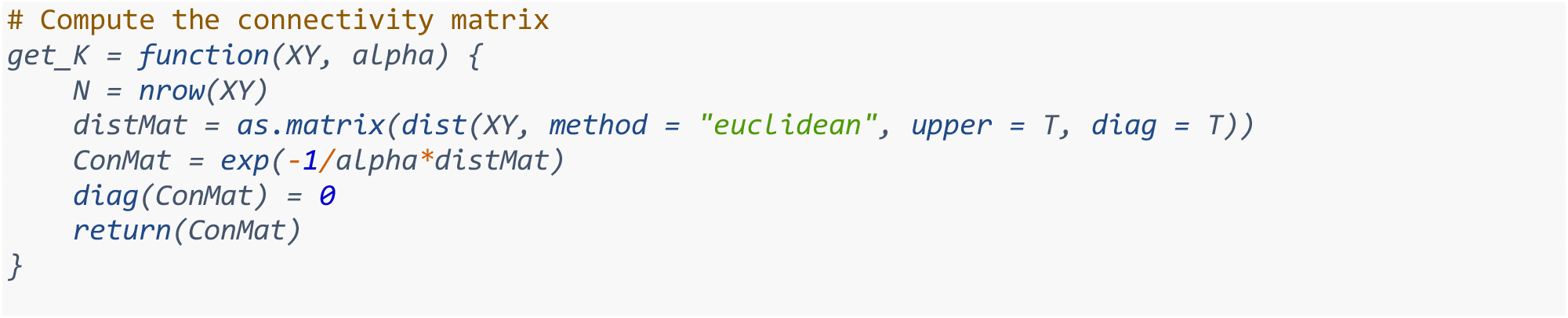

As an example, using the aggregated XY coordinates for 20 patches and our initial occupancy matrix with 8 species, we can see the connectivity between patches, and can calculate the contribution of immigration from each species to each patch.

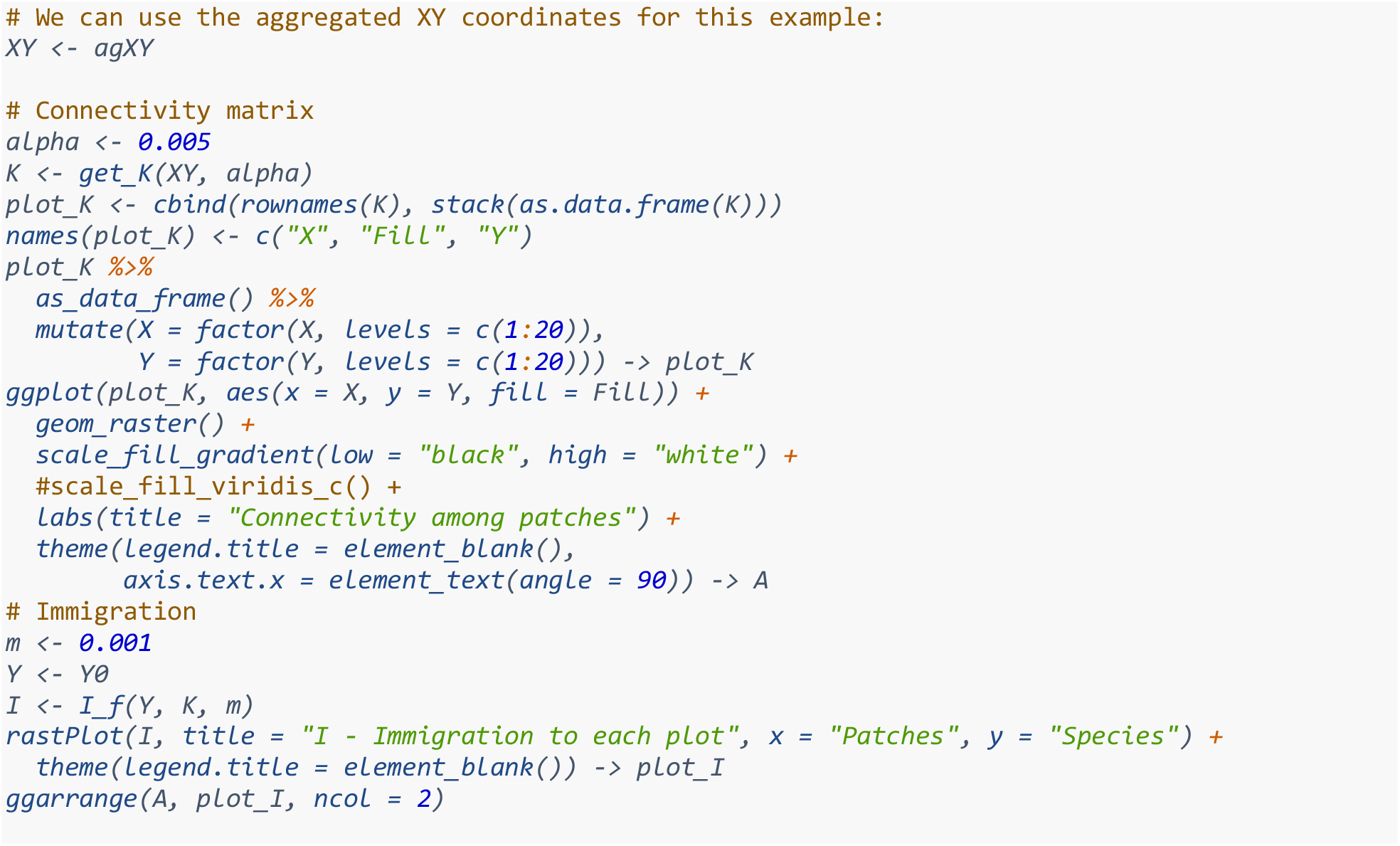

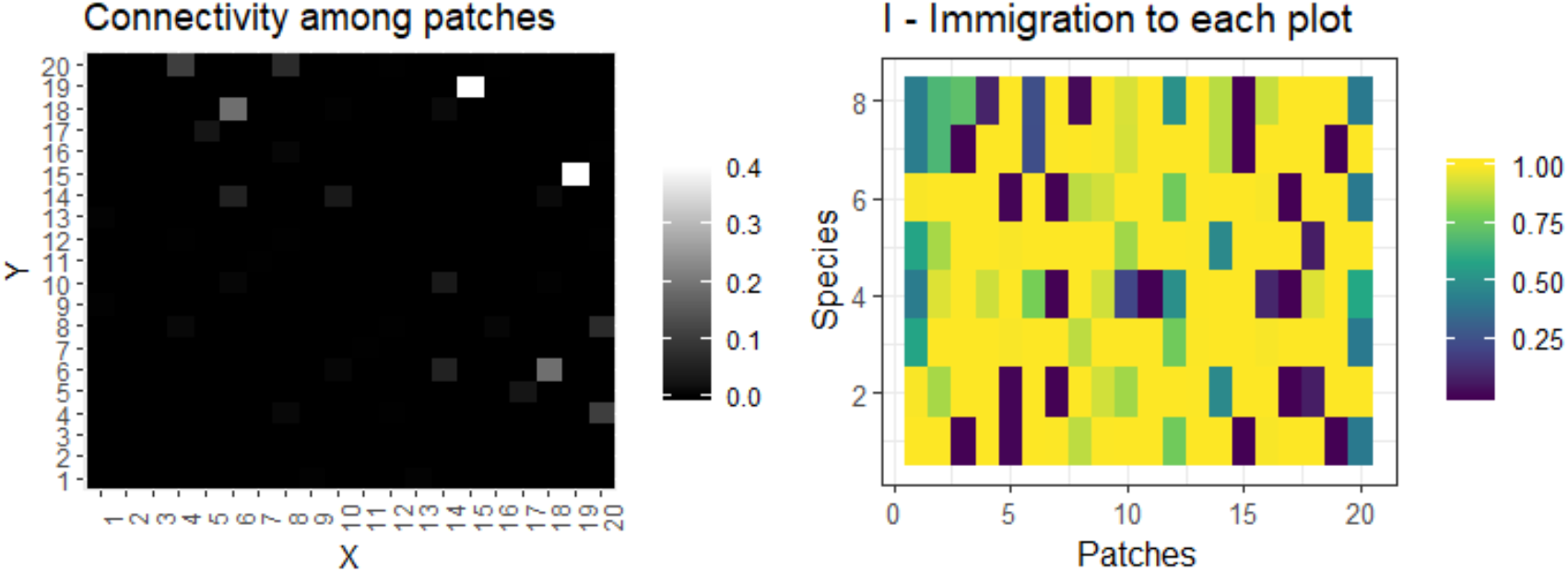

The effect of the environment on each species, depending on each species niche optima, is computed in the following code section. The argument `E` corresponds to the vector of values for the environmental variable in each patch. The other two arguments in this function are the niche optima (`u_s`) for each species and niche breadth(`s_c`).

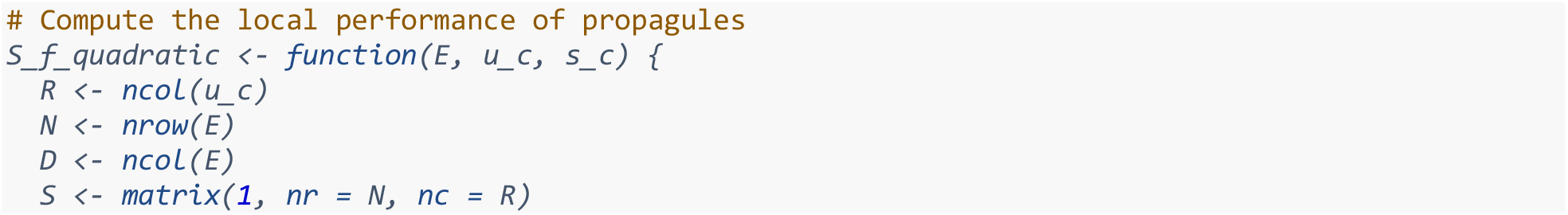

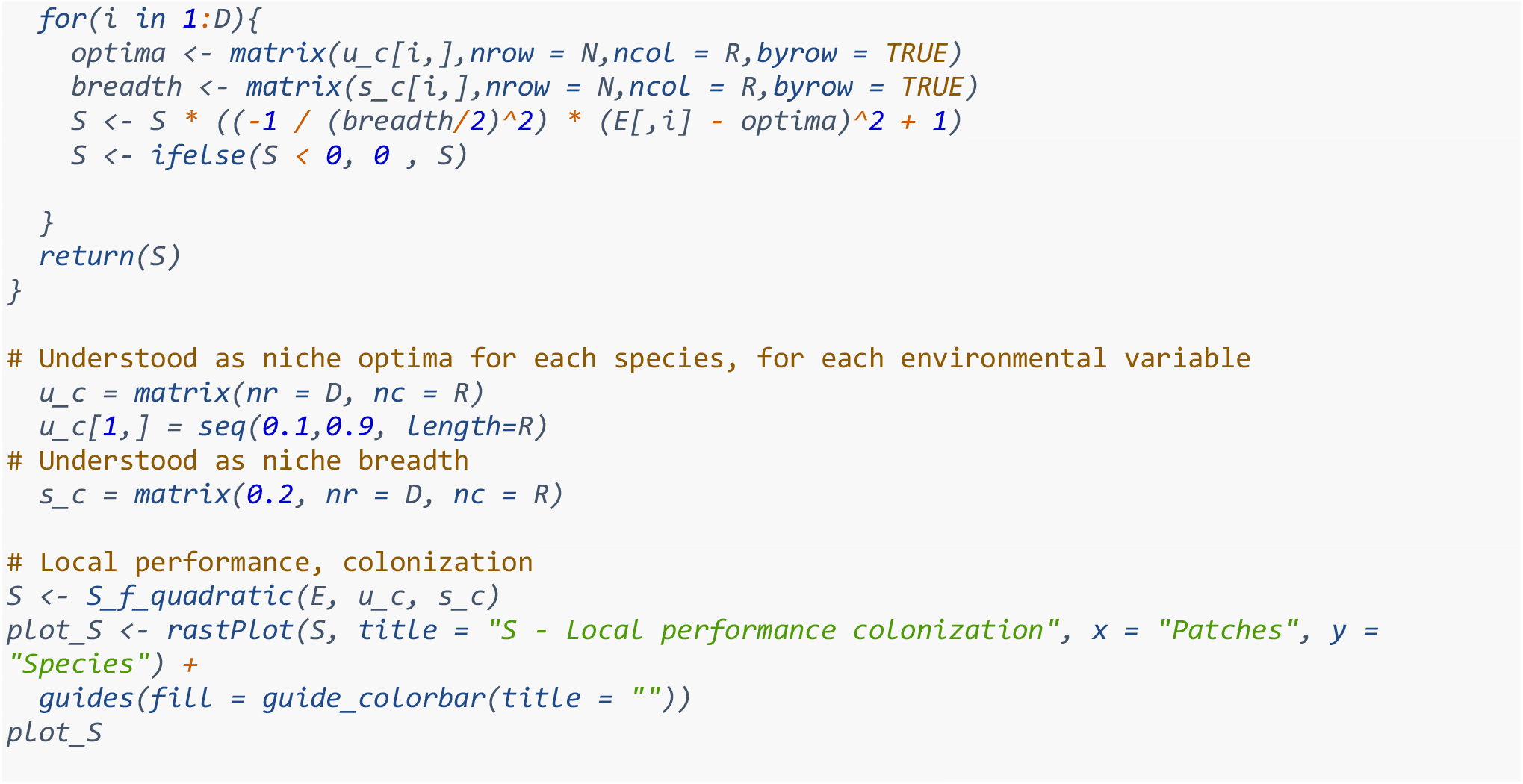

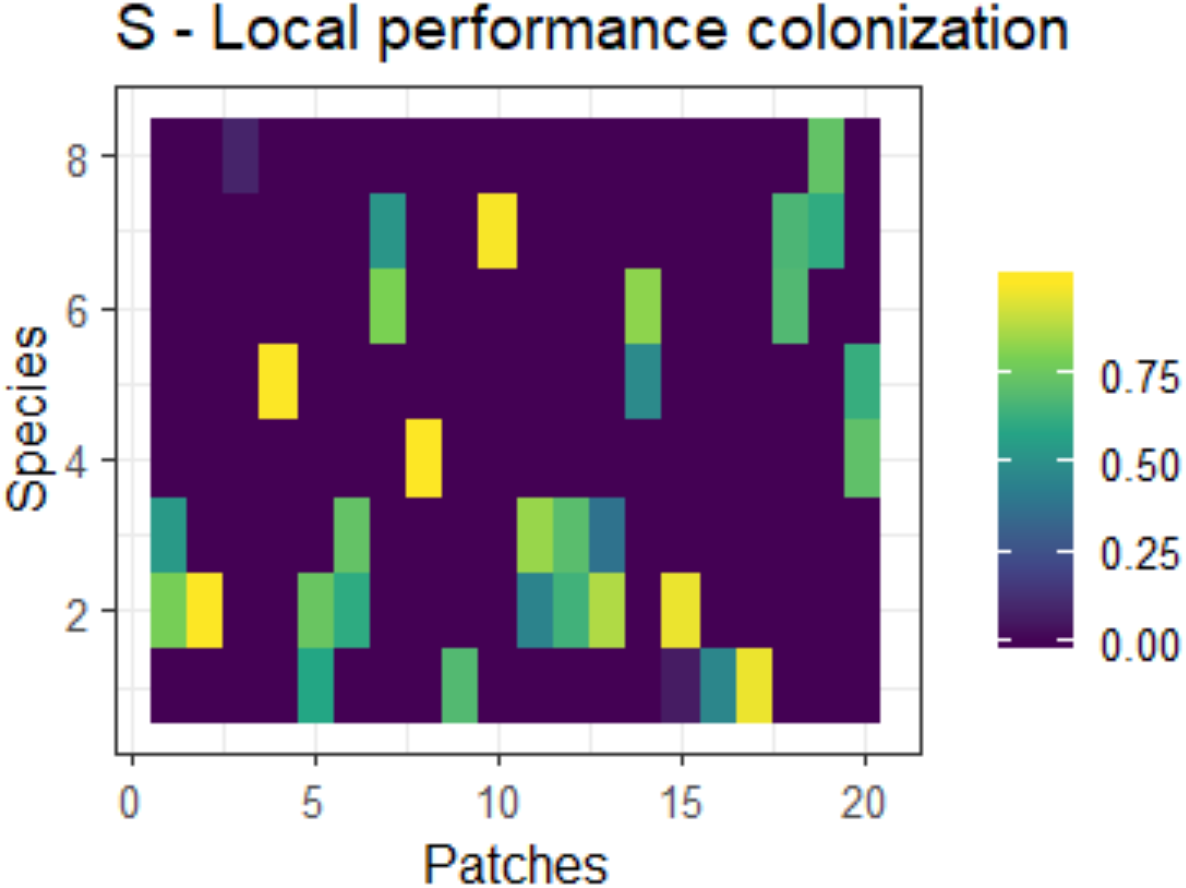

When incorporating species interactions into the toy model, we use the following interaction matrix *A*, where the colored black sections show species with potential of interacting:

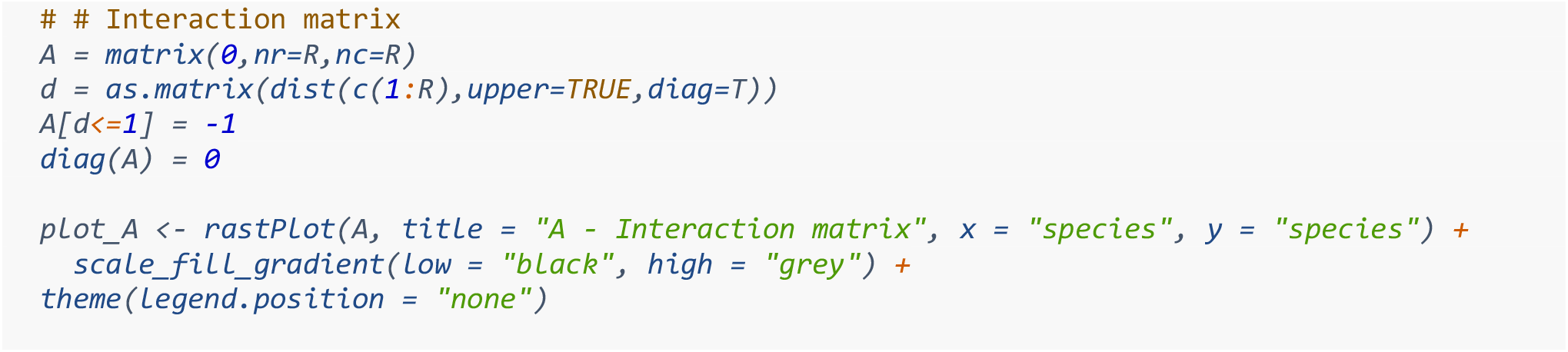

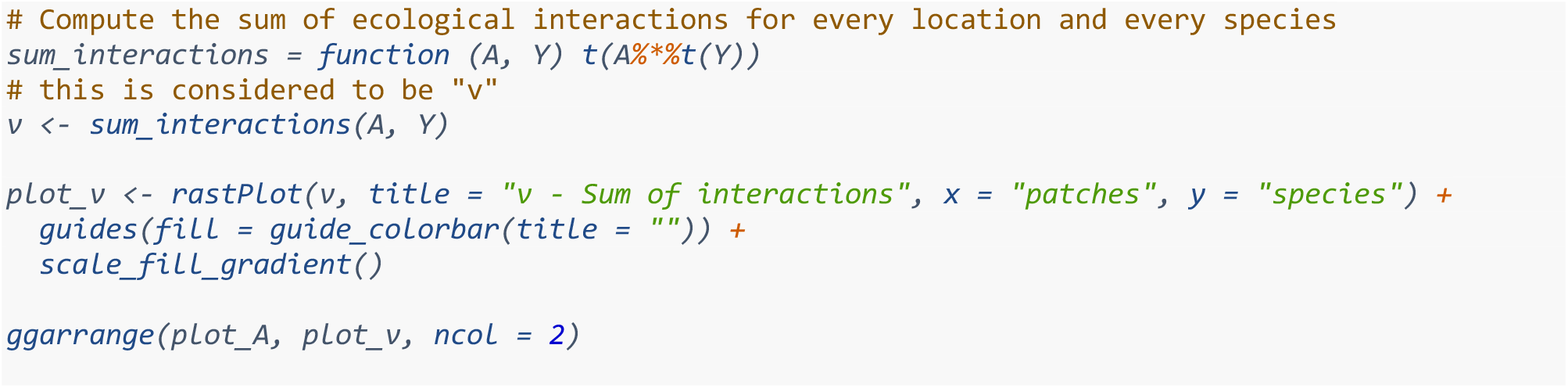

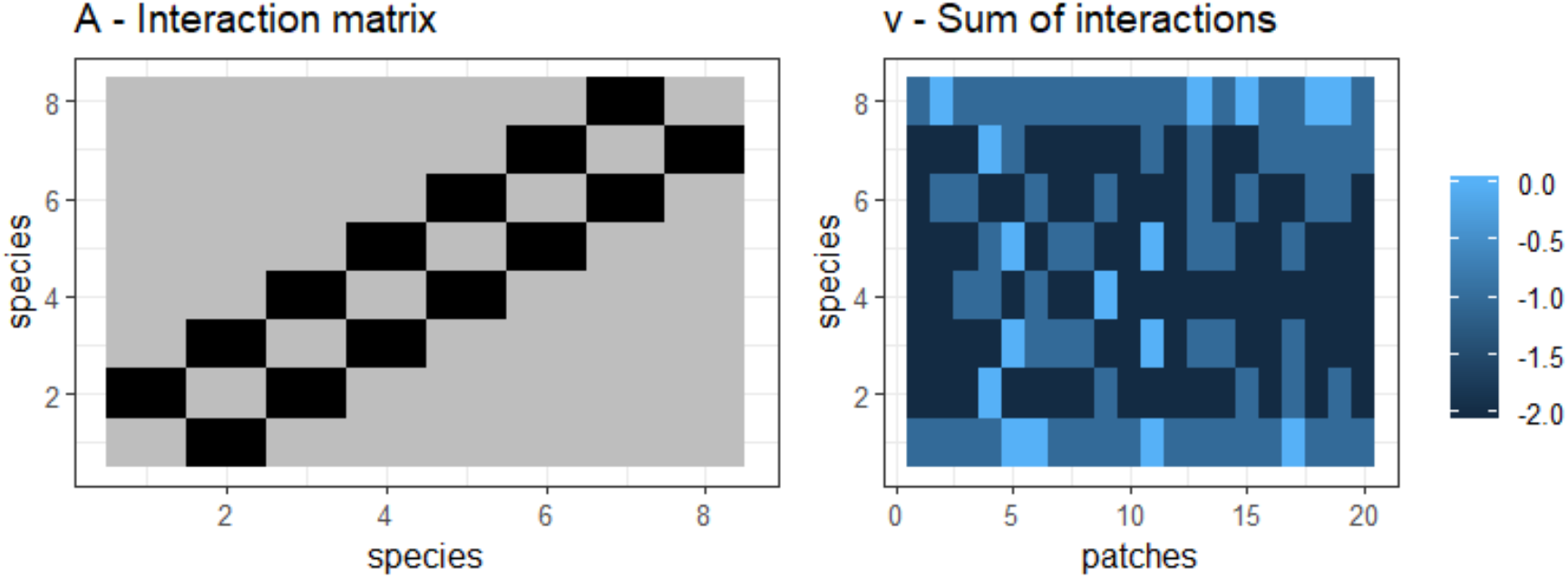

We compute the effect of ecological interactions on colonization probability with the function below.The arguments for this function are `v` as the resulting matrix from the sum of interactions, `d_c` as the sensitivity to interactions, `c_0` and `c_max` as the colonization parameters:

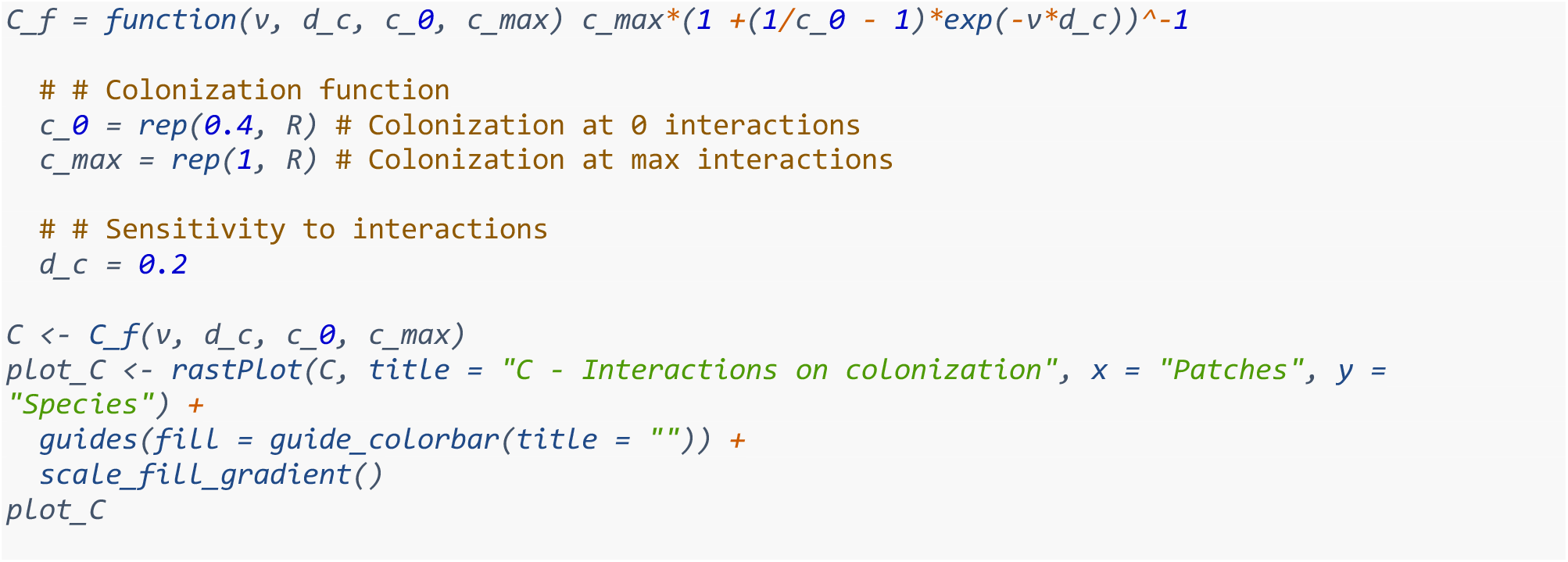

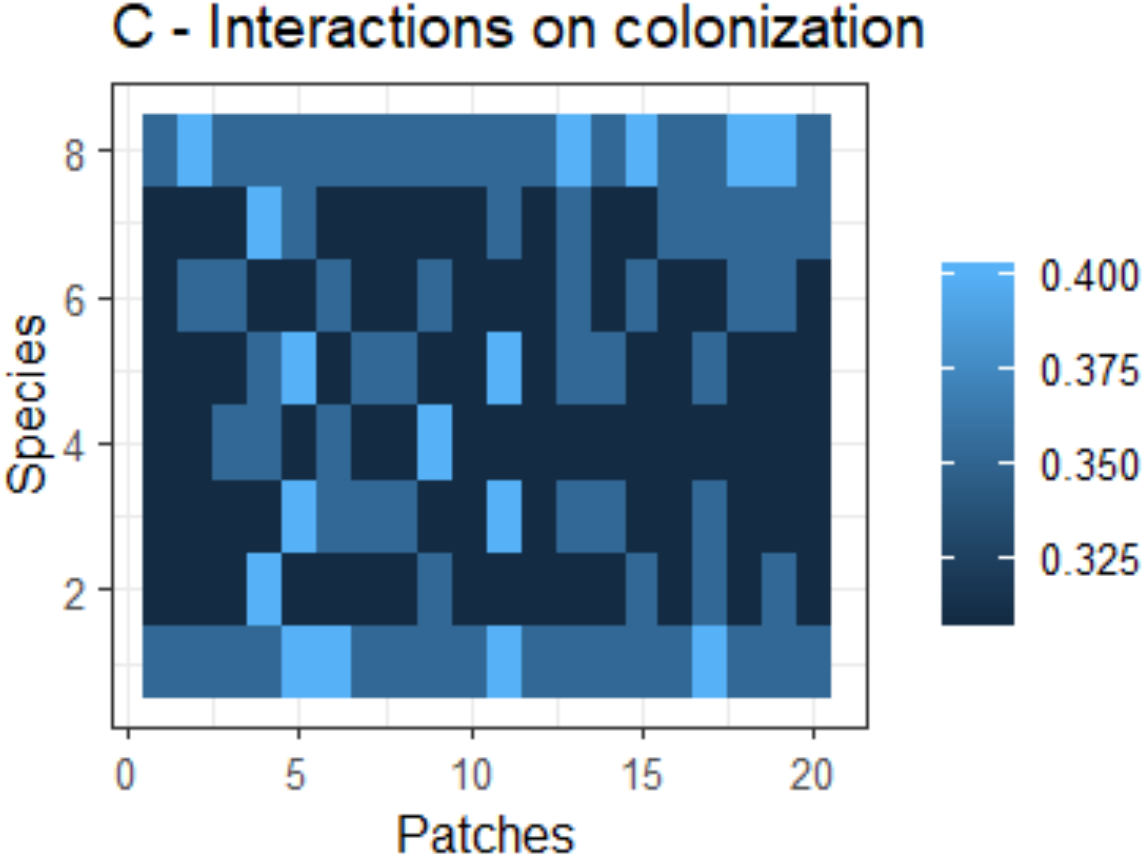

With all the components calculated, we can now compute the colonization probability

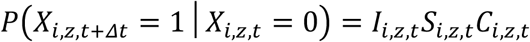

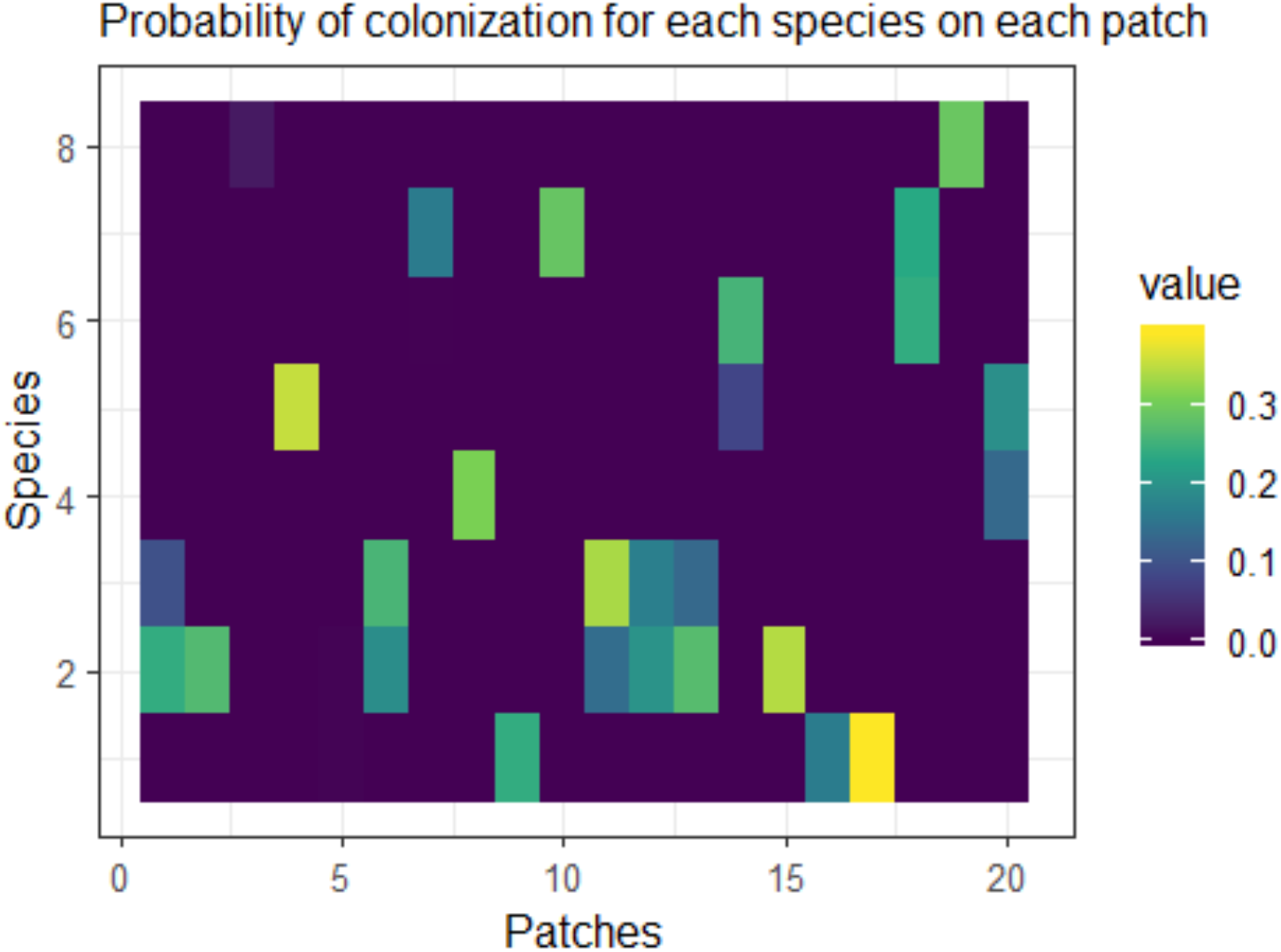

The following function calculates the effect of the environment on the extinction, with `E` being the environmental variable, and `u_e` and `u_s` being species level effect and the assymptote.

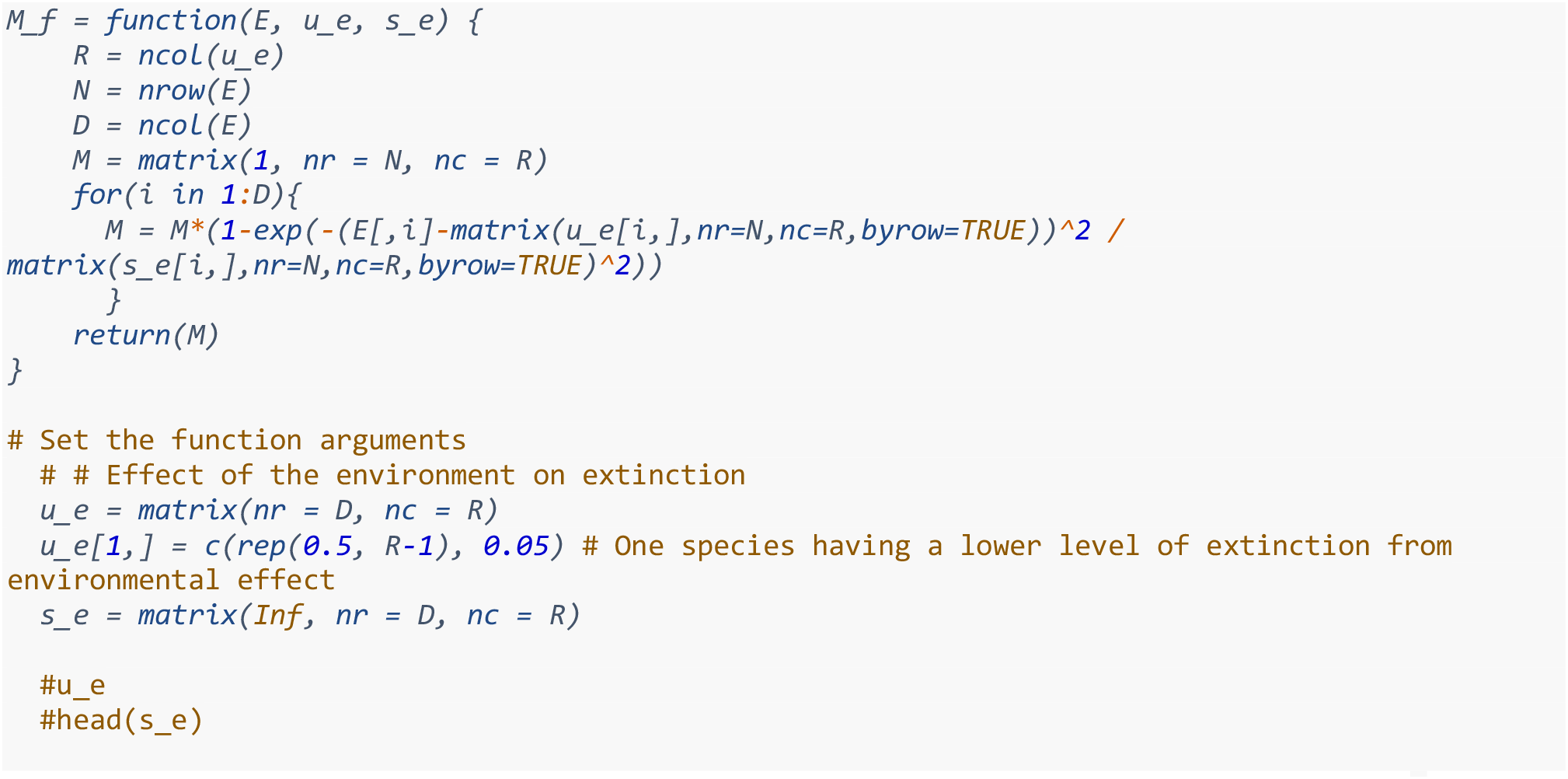

The following shows the effect of ecological interactions on extinction, using the same *v* matrix calculated above.

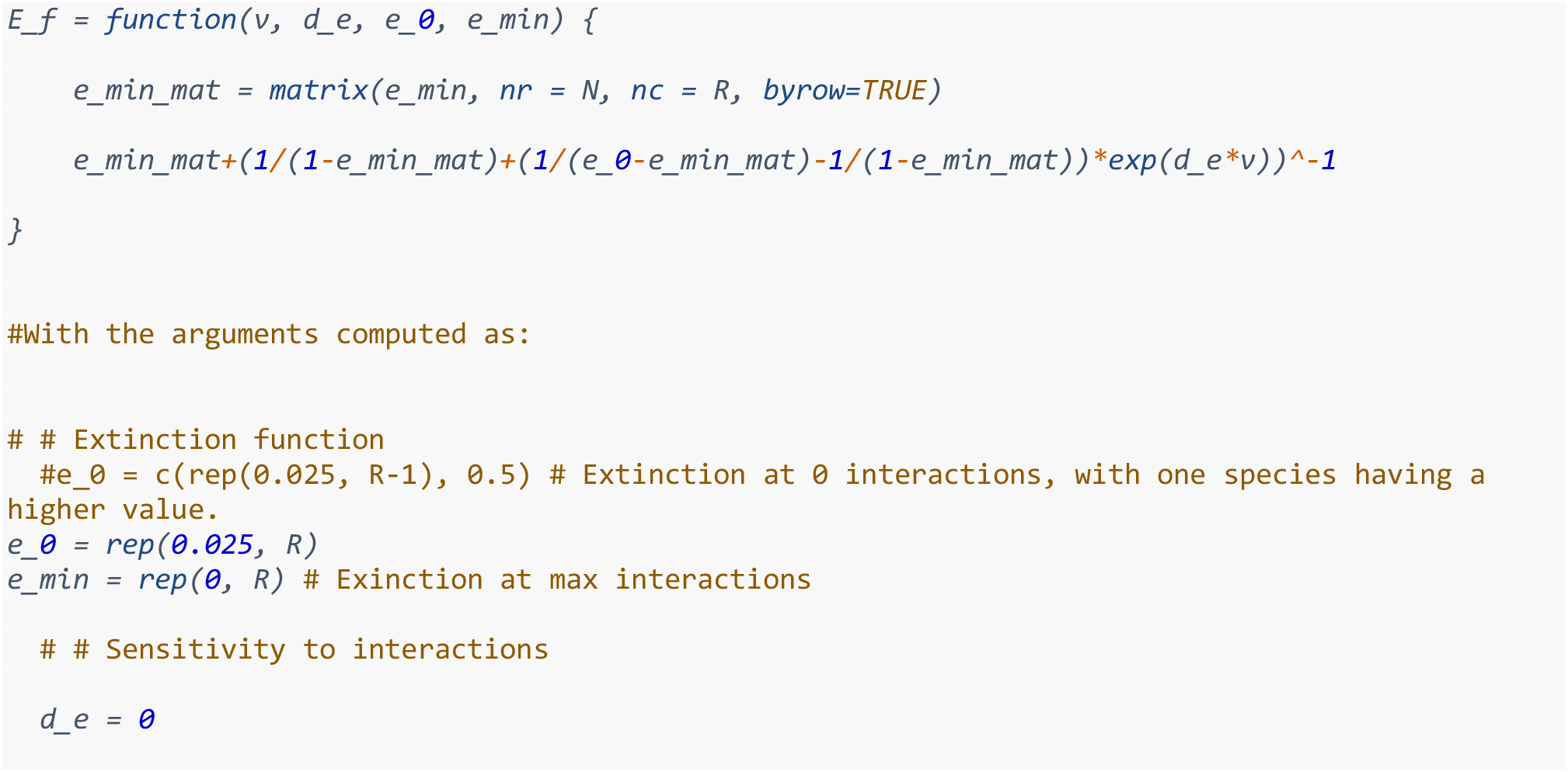

We can now compute the probability of extinction *P*(*X*_*i,z,t*+*Δt*_ = 1|*X*_*i,z,t*_ = 0) = *M*_*i,z,t*_*E*_*i,z,t*_ The figure shows these as identical graphs, since we have made all species have the same probability of extinction and the environment not having an effect on extinction.

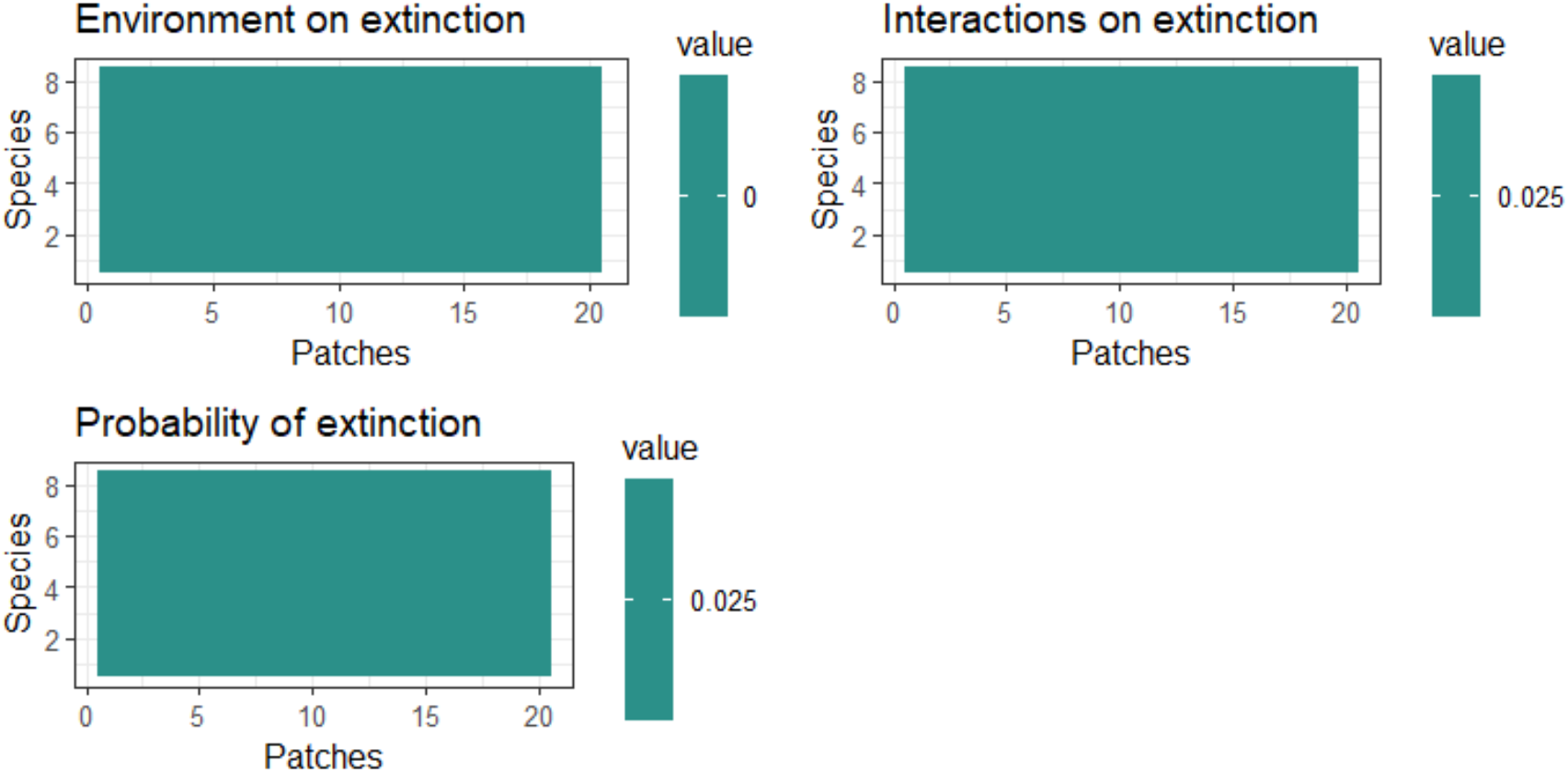

The way parameters are set, the extinction component is the same for all species in every patch.

#### Testing and changes

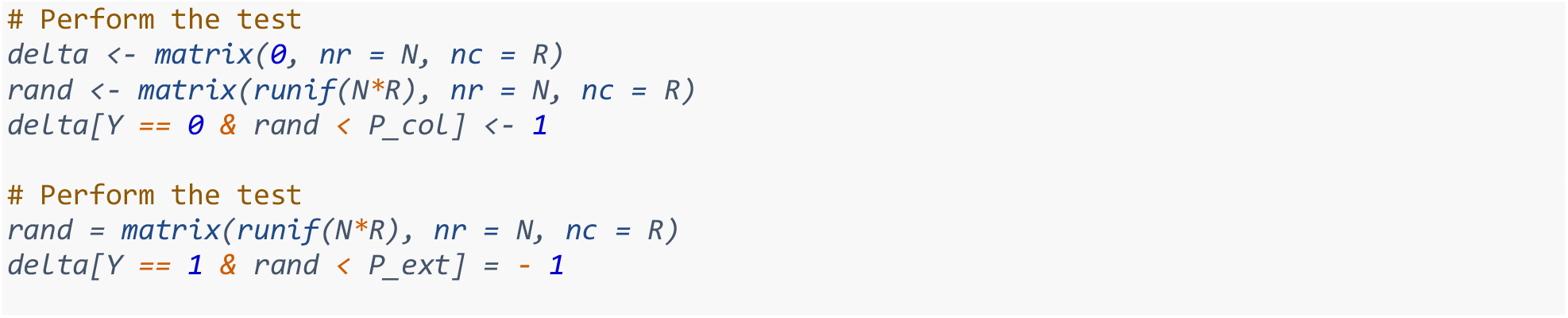

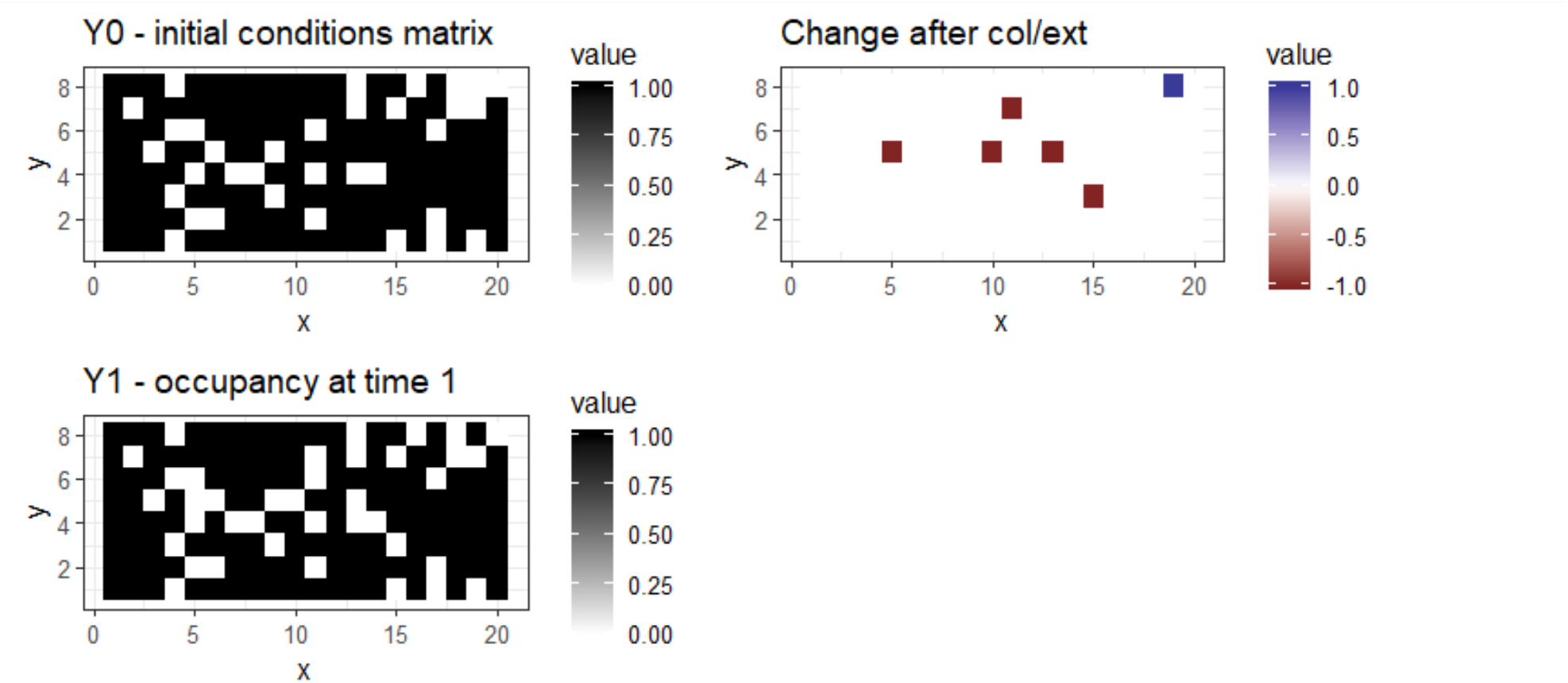

## Notes

### Competing Interest Statement

The authors have declared no competing interest.

